# Phased chromosome-scale genome assembly of an asexual, allopolyploid root-knot nematode reveals complex subgenomic structure: Analysis of the allopolyploid genome of *Meloidogyne javanica*

**DOI:** 10.1101/2023.03.20.533402

**Authors:** Michael R Winter, Adam P Taranto, Henok Zemene Yimer, Alison Coomer Blundell, Shahid Siddique, Valerie M Williamson, David H Lunt

## Abstract

We present the chromosome-scale genome of the allopolyploid root-knot nematode *Meloidogyne javanica*. We show that the *M. javanica* genome is predominantly allotetraploid, comprising two subgenomes, A and B, that most likely originated from hybridisation of two ancestral parental species. The assembly is annotated using full-length non-chimeric transcripts, comparison to reference databases, and *ab initio* prediction techniques, and the subgenomes are phased using ancestral k-mer spectral analysis. Subgenome B appears to show greater fission of chromosomal contigs, and while there is substantial synteny between subgenomes, we also identify regions lacking synteny that may have diverged in the ancestral genomes prior to or following hybridisation. Indels are common both between alleles within a subgenome, and between the A and B subgenomes, suggesting the *M. javanica* genome exists in a dynamic hypo-tetraploidy where copy number can vary along the chromosome. This annotated and phased genome assembly forms a significant resource for understanding the origins and genetics of these globally important plant pathogens.

**Author Summary:** Root-knot nematodes represent one of the most significant crop parasites globally. Despite their agricultural importance, only rudimentary genomic resources have been published to date, leaving a gap in the understanding of genetic mechanisms driving genome evolution and crop virulence. Here, we have used modern genomic and bioinformatic approaches to create a chromosome-scale reference genome to investigate the origins and genomic constitution of the root-knot nematode species *Meloidogyne javanica*. This species reproduces by ameiotic parthenogenesis and has an allopolyploid genome and is among the most damaging plant parasitic nematodes with a large and evolving plant host range.

Utilising modern long-range DNA sequencing and bioinformatics approaches, we successfully phased the genome into its constituent subgenomes, a first for this agriculturally important clade. While we find the genomic landscape is mostly syntenic between subgenomes, we identified regions of minimal similarity, and highlight structural divergence between subgenomes. We demonstrate that this species was originally tetraploid, but insertions and deletions have been the major force in generating diversity, resulting in a hypo-tetraploid genome with local variations in ploidy.

## Introduction

### The assembly of allopolyploid genomes

Allopolyploidy is a genomic state characterised by more than two chromosomal complements, with one or more of these complements resulting from a hybridisation event leading to the presence of distinct (homoeologous) subgenomes within a single cell <u>[1]</u>. Allopolyploids may account for 11% of plant species including many model species and important crops <u>[2]</u>. Although not as frequent as in plants, genomic investigations are indicating that ancestral genome duplication, hybridisation, and complex genome arrangements are more widespread than previously recognized in animals <u>[3,4]</u>. Assembly and analysis of allopolyploid genomes, however, is challenging for a number of reasons <u>[5]</u>. The increased number of alleles within an allopolyploid genome can interfere with algorithms used by many assemblers leading to the accumulation of switch errors; regions of the assembly where the sequence switches between haplotypes or homoeologs. In addition, the high amount of repeat content often found in allopolyploid genomes can result in fragmentation of the final assembly if sequenced reads fail to span the repeat <u>[6,7]</u>. Another difficulty in accurate assembly of allopolyploid genomes has been ‘phasing’ ie, the assignment of assembly contigs to the correct subgenome. Switch errors and misassemblies introduced during the assembly process can impair the signals required to successfully phase a scaffold, and potential crossover interactions between homoeologs can further complicate this signal <u>[8,9]</u>.

Assembly of allopolyploid genomes has become more feasible due to the advent of long-read sequencing technologies and better assembly algorithms. Most chromosome-scale allopolyploid assemblies in the literature are of agricultural plants <u>[10–13]</u>, although a few chromosome-scale allopolyploid assemblies of animal genomes are now also available <u>[14,15]</u>.

### Root-knot nematodes

Root-knot nematodes (RKN) - genus Meloidogyne - are a group of obligate plant parasites that include species which severely reduce crop yield [16]. Second-stage juveniles (J2s) of RKNs hatch in the soil and are non-feeding, needing to invade a host plant root to complete their life cycle. Upon reaching the vascular cylinder, J2s induce the formation of a feeding site inside the root, characterised by formation of a gall (“root-knot”) and highly modified “giant cells” on which the nematode feeds [17]. Three closely related species within the Meloidogyne genus, M. arenaria, M. incognita, and M. javanica, which we refer to here as the Meloidogyne incognita group (MIG), have extremely broad host ranges spanning the majority of flowering plants [18,19]) and together are estimated to cost the agricultural industry tens of billions of US dollars a year [20–22]. Almost 100 RKN species have been described [23,24], which differ in host range, pathogenicity, geographic range, morphology and reproductive mode.

Although RKN nematodes species have diverse modes of reproduction including amphimixis, automixis, and obligate apomixis, cytological examination indicates that M. javanica and most other MIG species reproduce by mitotic parthenogenesis; that is, maturation of oocytes consists of a single mitotic division in which chromosomes remain univalent at metaphase [25,26]. Phylogenomic analysis has revealed that each species possesses two divergent copies of many genes, that the three species likely originated from interspecific hybridisation, and that they share the same ancestors who have provided the A and B subgenomes [19,27,28].

Despite asexual reproduction, field isolates and greenhouse selections of MIG species are diverse and successful, differing in their ability to reproduce on specific crop species and varieties (Hartman and Sasser, 1985; [20,29–31]. A widely investigated example is acquisition of ability to reproduce on tomato with the resistance gene Mi-1, which confers effective resistance to MIG species and is widely deployed for nematode management in tomato [32]. Many independent studies have identified MIG populations that are able to break Mi-mediated resistance; these include both field isolates and greenhouse selections of isofemale lines [33,34]. However, efforts to decipher the genetic mechanisms for these phenotypic variants have not so far been successful due in part to the lack of tractable genetics and limitations in genome assemblies. Current MIG genome assemblies are fragmented and the homoeologous genomes are mostly unphased [19,28,35,36] making it difficult to compare homoeologous sequences, gain a true picture of diversity, or to understand the nature of functional variation.

Here we apply a combination of modern genomic and bioinformatic approaches to generate a highly-contiguous, chromosome-level assembly of M. javanica, phased into two subgenomes, creating the first chromosome-scale genome assembly of an apomictic allopolyploid animal that we are aware of. This assembly should provide a very valuable framework for research into the diversity and functional divergence of plant pathogenic nematode species. In the wider research landscape, genomes such as those from within the MIG can aid in our understanding of adaptation, ploidy, and evolution of genomes following hybridisation events and loss of meiosis [37].

## Results

### Sequencing and profiling of read libraries

We used PacBio single-molecule real-time (SMRT) sequencing technology, Hi-C chromatin conformation capture, Nanopore long-read sequencing, and Iso-Seq RNA sequencing to generate a genome assembly of *Meloidogyne javanica* strain VW4. Following quality control and concatenation of two libraries, we obtained 2,255,922 PacBio HiFi reads totalling 35.39 gbp (ACC: XXXXXX; S1 Fig). After quality control and concatenation of Oxford Nanopore PromethION data, we obtained 340,373 reads totalling 17.52 gbp (ACC: XXXXXX) (S2 Fig). Our Hi-C library contained 375,330,537 read pairs, of which 26.20% were of sufficiently high quality for scaffolding with *Proximo* (Phase Genomics, WA) (ACC: XXXXXX). After demultiplexing of our Iso-Seq library, we obtained 2,506,897 full-length non-chimeric sequences which collapsed into 59,637 high quality isoforms (ACC: XXXXXX).

Genome-wide k-mer profiling of concatenated PacBio HiFi read libraries with *smudgeplot* <u>[38]</u> indicated that 48% of the genome was tetraploid, 22% was triploid, and 29% was diploid (Fig 1A). However, this analysis likely underestimates the proportion of tetraploid regions, as crossover events or conversion between subgenomic copies can cause homogenisation. An incomplete or hypo-tetraploid state was also indicated using a k-mer spectra approach by *GenomeScope2*, predicting a haploid genome length of 68 mbp, and a duplication rate of 3.4 <u>[38]</u> (Fig. 1B). This duplication rate is similar to the value seen for CEGMA genes in other assemblies of *M. javanica* (3.68) <u>[28]</u>.

**Fig 1:**
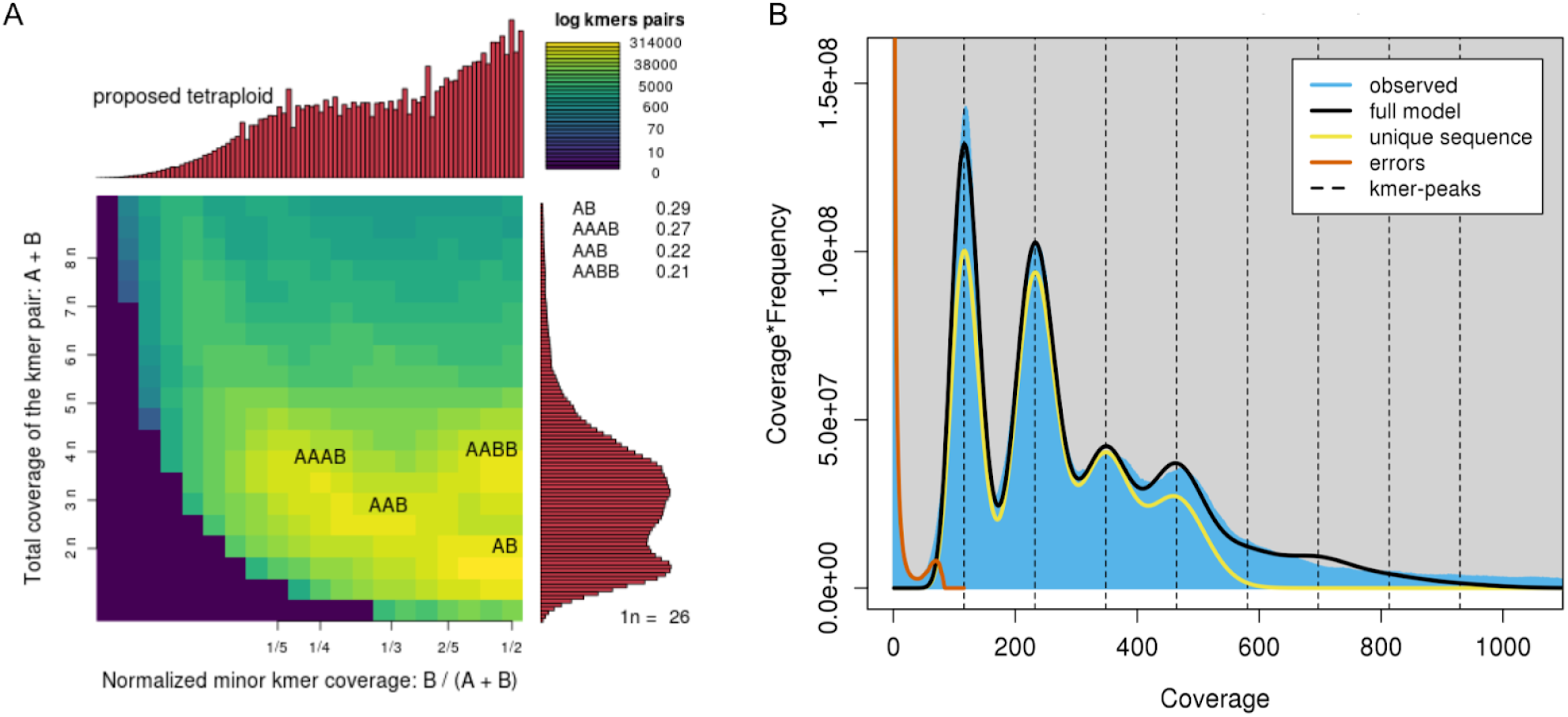
Genome profiling plots. (A) Smudgeplot (left) proposing M. javanica as tetraploid, reporting the predicted percentages of ploidy levels in the genome as follows: tetraploid (48%), triploid (22%), or diploid (29%). (B) GenomeScope2 plot (right) showing four distinct peaks in both the predicted model of tetraploidy (black line) and in the observed k-mer spectra (blue fill). The amount of unique sequence falls to zero shortly after the fourth peak, indicating that k-mers at higher ploidies than four were mostly repetitive elements.

### Assembly and annotation

#### Scaffolding and final assembly

From our draft assemblies of the PacBio data, we carried forward an iteration assembled using *HiFiasm* <u>[39]</u> based on overall contiguity and comparison to expected diploid genome length. Following purging of duplicates from the PacBio assembly and scaffolding with Oxford Nanopore reads, we obtained 37 contigs with the N50 of 9.5 mbp. Hi-C scaffolding with the *Proximo* pipeline identified 16 chromosome level clusters (Phase Genomics Ltd), but increased the total number of scaffolds from 37 to 66. Following scaffolding with Hi-C, *samba* <u>[40]</u> joined some small scaffolds and fragmented the largest scaffold in the assembly, increasing the number of scaffolds from 66 to 69.

The final assembly scaffolds contains 150,545,692 bp with an N50 of 5,793,182 bp, at 30.11% GC content, and overall, 99.96% of reads in our combined PacBio HiFi library map successfully back to the assembly, indicating a high level of completeness (S1 Table). Of the total assembly, 97.87% was contained in the longest 33 scaffolds (S3 Fig), which ranged from 898 kbp to 9,595 kbp in length (Fig 2A; S6 Table). These 33 scaffolds contained 99.85% of all transcribed gene models. Of the remaining 36 scaffolds, 4 were identified by *blobtools* <u>[41]</u> as likely contaminants (Arthropoda, Chordata, and Streptophyta; S4 Fig). One further contig was the *M. javanica* mitochondrial genome (Scaffold 67). The remaining 31 small contigs were all less than 215 kbp, with a mean length of 88.9 kbp, and since they contained few identifiably functional elements (0.19% of gene models), we excluded them from the final analysis.

**Fig 2.**
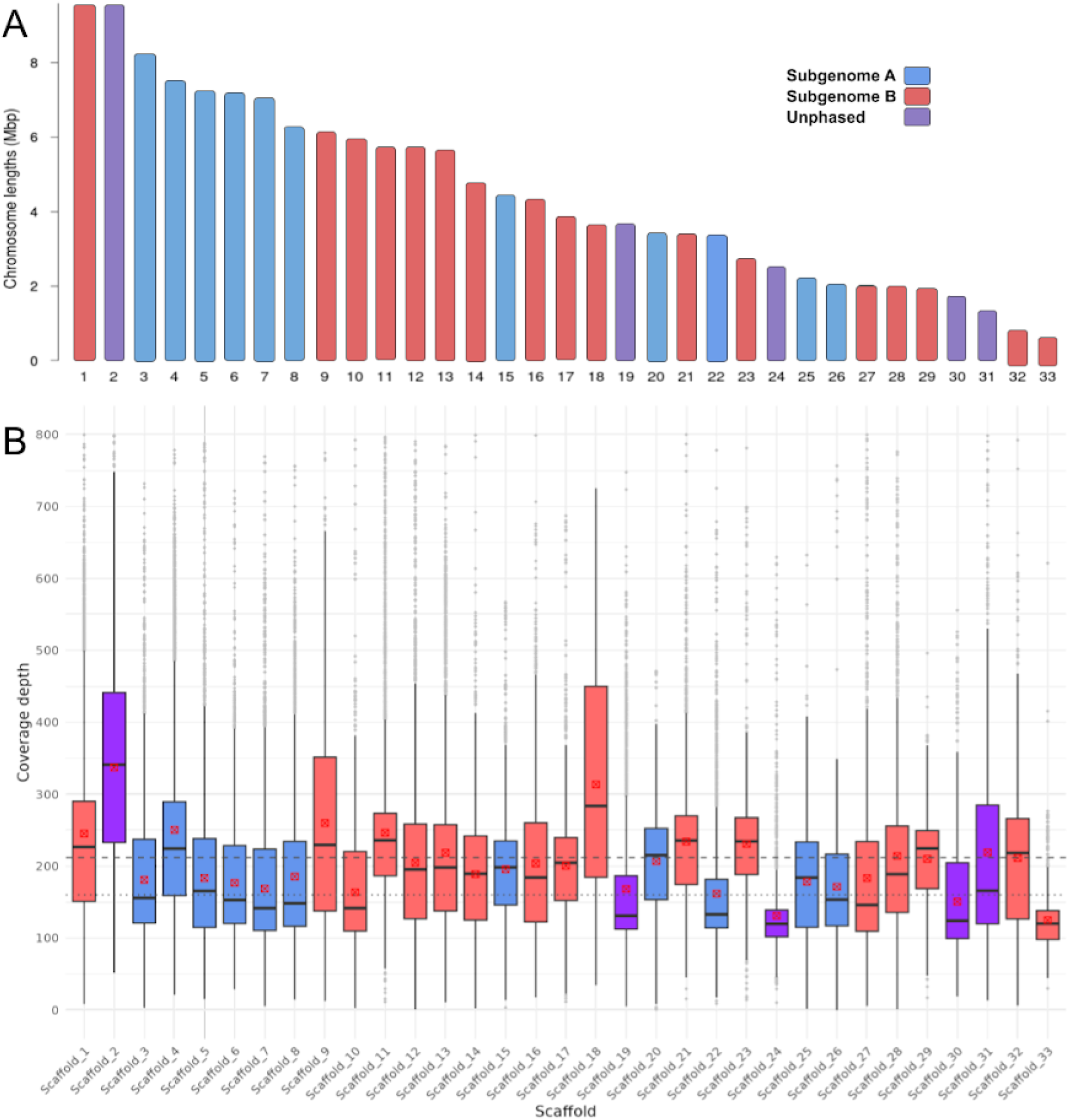
Ideogram and coverage depth of longest 33 scaffolds. Scaffolds are coloured according to phasing status; blue - subgenome A, red - subgenome B, purple - unphased. A, Ideogram of 33 largest scaffolds. These 33 scaffolds contain 98% the total length of the assembly, with the remaining 36 contigs being shorter than 250 kbp, containing few gene models (0.19%), and consisting of mostly repetitive elements. B, Boxplot displaying the distribution of coverage depth for each scaffold. Red points denote the mean of data in each box. Coverage has been limited to a maximum of 800x to exclude probable repetitive sites with anomalous coverage depth. Dashed line shows the overall mean for all coverage levels across 33 scaffolds. Dotted line shows the mode of all coverage across 33 scaffolds.

#### Repeat annotation

In total 30.46% of our assembly was identified as repetitive elements, with 4.94% identified as retroelements, 3.77% as DNA transposons, while 16.88% remain unclassified repeats (S2 Table). A total of 164,394 transcripts, representing 59,632 isoforms, were detected through mapping of our Iso-Seq library. *MAKER3* detected a total of 22,433 genes, containing 227,617 exons, 10,044 5’ and 5,253 3’ UTRs (S3 Table).

#### Core Eukaryotic Gene and Single Universal Copy Ortholog Analysis

Analysis of the final assembly with *CEGMA<u>[42]</u>* detected 233 of 248 CEGMA genes (93.95%). This is a higher level of completeness than the most recent *M. javanica* assembly, and comparable with contemporary *Meloidogyne* assemblies <u>[19,36,43,44]</u>

The average number of orthologs for each complete CEGMA gene (a proxy for ploidy) is 1.88 indicating that only ∼6% of the diploid genome is unassembled or not present in the biological chromosomes. 173 complete BUSCO genes were detected, representing 67.9% of genes in the BUSCO eukaryote database. Of the BUSCOs identified, 31.4% were duplicated (S5 Fig).

### Coverage and ploidy analysis

Mean coverage depth for all scaffolds - excluding mitochondrial - was 206.6x, falling to 206.1x for the longest 33 scaffolds (Fig 2B). For some scaffolds (2, 9 and 18) coverage depth was significantly in excess of this average, indicating collapsed regions; that is, representation of three or four rather than two copies. Collapse is expected to arise in a polyploid assembly when homoeologous sequences are similar enough at a nucleotide level that they are inferred to be multiple homologous alleles of the same region, which are then collapsed to form the reference assembly copy <u>[8]</u>. Other scaffolds (19, 22, 24, 33) were underrepresented suggesting that they may be present as a single copy.

We then examined the coverage depth frequency distribution for each scaffold (Sup Fig 7). For phased scaffolds representing two identical homologs, the frequency of coverage for all bases in each scaffold would be expected to have a single peak (240x). However, while predominantly single peaks were seen for under-represented scaffolds thought to be present as a single copy (19, 22, 24, 33), we observed two peaks for the majority of scaffolds (Fig 3A & C; S7 Fig). Two peaks would be expected if homologs are not identical and similarity is disrupted by indels, with the second peak representing the reduced coverage of the hemizygous region (∼120x). For scaffold 2, which is over-represented in coverage depth, four clear peaks are present (Fig 3) indicating that more than two copies map to the scaffold. This likely resulted from exclusion of the scaffold’s homoeolog from the assembly, leading to an assembly collapse as noted above. Nevertheless, the presence of 4 peaks is consistent with the presence of polymorphisms between homologous copies as well as between homoeologous pairs.

**Fig 3.**
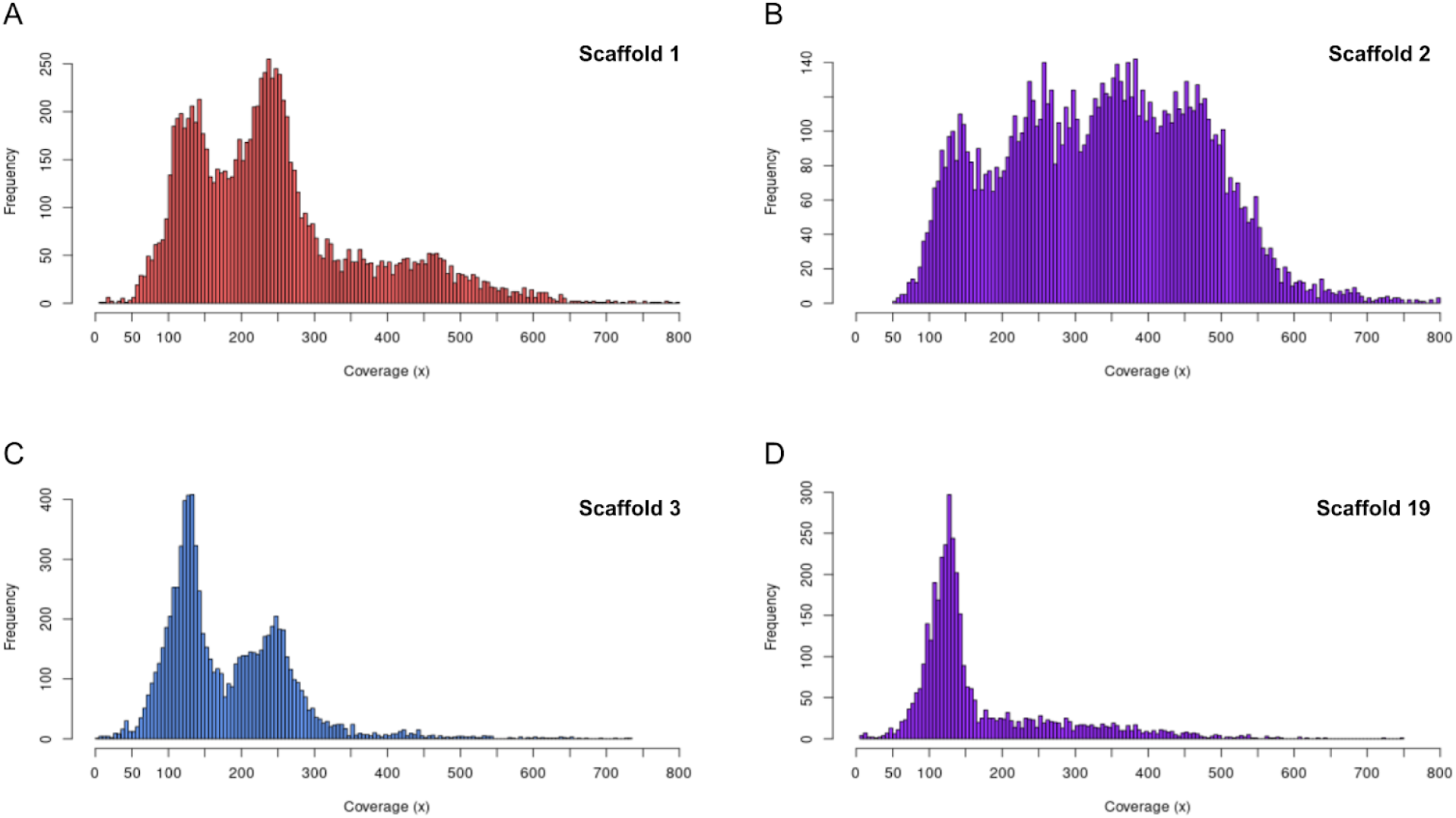
Coverage depth frequency distributions of scaffolds 1, 2, 3, and 19. X-axis represents coverage depth and y-axis represents frequencies of coverage. Colour indicates phase status: red for subgenome B, blue for subgenome A, and purple for unphased scaffolds. (A) Scaffold 1, top left, displays two main peaks of coverage with a small tail suggesting short collapsed regions. (B) Scaffold 2, top right, displays four peaks of coverage, indicating that much of this scaffold is collapsed and four copies are mapping to it. (C) Scaffold 3, bottom left, shows two peaks and very little tail, indicating two copies mapping and little to no assembly collapse. (D) Scaffold 19, bottom right, shows only one peak at ∼120x coverage, suggesting that only one copy maps to this scaffold.

To examine the ploidy distribution in another way, we plotted a sliding window of coverage across each scaffold (S8 Fig). In support of the coverage depth distributions, the most frequent outcome was that scaffolds in the assembly show two layers of stratification in coverage at a constant proportional depth, indicating two copies distinguished by indels between them. This pattern was strongest for scaffolds with two coverage peaks, particularly phased homoeologs (discussed below, Fig. 3 A & B; S7 Fig). Scaffold 2 and scaffold 18, which are over-represented in sequence depth, contain regions of four levels of coverage depth covering much of the length of both (Fig 3B; S8a and 8d Figs). This increased amount of stratification at proportionally higher coverage depths (360x and 480x) reveals assembly collapse, where four copies map to a single site. Together these coverage depth results suggest that the *M. javanica* assembly represents two homoeologous subgenomes (∼87% of total length), with only 13% of the assembly unphased or collapsed.

### Identification of homoeologous pairs and phasing of subgenomes

Given the diploid nature of the assembly, which represents each subgenome as a single copy, we expected to find scaffolds from these subgenomes present in homoeologous pairs. Through detection of shared orthologs (Supplementary Methods) twenty pairings were identified between the longest 33 scaffolds of the assembly, each sharing between 20 and 351 CDS orthologs (Fig 4, S4 Table). Alternative methods of identifying homoeologous pairs were corroborative (S5 Table). Some pairs were not mutually exclusive, and exhibited CDS links to scaffolds outside of the primary pair, suggesting translocation and syntenic changes between them. All scaffolds that showed two depths of coverage were assigned as homoeologous pairs as expected if each subgenome was heterozygous. Four scaffolds were excluded from pairings including scaffold 2 which showed high amounts of collapse and scaffolds 19, 24, 30, and 31 which are relatively small and/or present as single copies (S7 Fig and 9; S6 Table).

**Fig 4.**
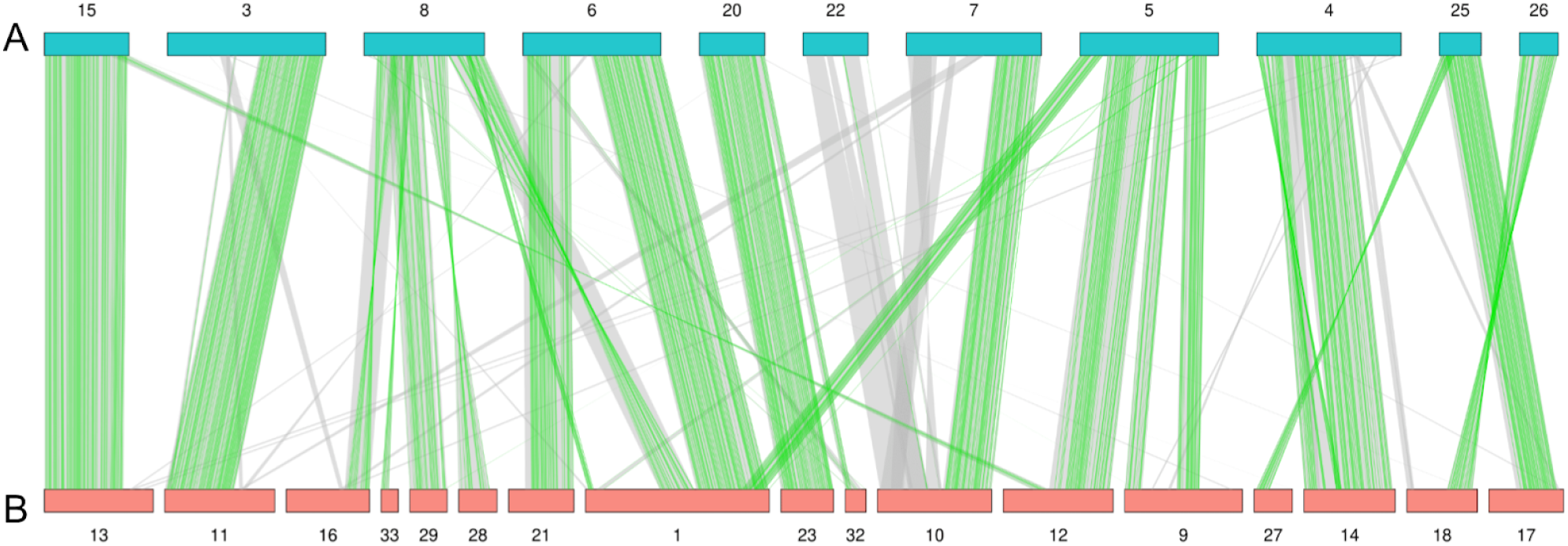
Macrosynteny analysis between subgenomes. Glyphs along top and bottom represent scaffolds assigned to either subgenome A (blue) or subgenome B (red). Green lines mark locations of synteny between transcribed genes identified from mapping of Iso-Seq sequences. Grey lines mark locations of synteny between genes identified through *MAKER3* gene prediction. Synteny and collinearity were identified using the *MCScan* module of *JCVI* using Iso-Seq informed transcriptional annotation. Scaffolds that could not be assigned to a subgenome are not shown.

In order to assign contigs to a subgenome, we used a modified version of an ancestral k-mer spectra analysis <u>[45]</u> (Supplementary Methods). This approach is based on the premise that the allopolyploid’s subgenomes possess repeats that diverged in the two parental genomes before hybridisation. This results in a distinguishable signature in each subgenome’s k-mer spectrum, and allows us to determine the parental species from which a given sequence descends. We successfully phased 85.39% of the genome into A or B subgenomes (Fig 2A; S9 Fig; S6 Table). Scaffold 2, which exhibits extensive assembly collapse, did not phase using these methods. Some smaller scaffolds that did not phase by k-mer based methods were later assigned to a subgenome based on the phase status of their opposing homoeologous scaffold (S6 Table). Of the total length of the final assembly, 39.15% was assigned to subgenome A and 46.24% was assigned to subgenome B, leaving only 14.61% unassigned.

### Subgenomic synteny analysis

Comparison of annotation of scaffold pairs assigned to subgenomes A and B revealed both long regions of synteny and large structural differences between subgenomes (Fig 4). Scaffold 13 and scaffold 15 are syntenic along almost the entire length of the shorter homoeolog and share high nucleotide similarity throughout. Similarly, scaffold pairs 20 and 23, as well as 17 and 25, share long syntenic blocks of shared CDS and long runs of high nucleotide similarity.

Scaffolds 7 and 10 are syntenic for almost half of their length whilst the remaining sequence lengths share no synteny and have very low nucleotide similarity. Synteny analysis and scaffold comparison suggest that chromosomal fragmentation has occurred. For example, scaffold 8 of subgenome A shows long collinear blocks with scaffolds 16, 28, 29, and 33 of subgenome B. Re-examination of Hi-C and ONT long-read scaffolding as well as manual inspection of read mapping support this fragmentation. Scaffold 1 of subgenome B shares syntenic blocks with scaffolds 5, 6, and 8 of subgenome A, in an order that suggests chromosomal structural differences between the subgenomes. Many phased scaffolds also exhibit small amounts of extra-pair synteny indicating numerous small translocations throughout the genome.

## Discussion

### Assembly of the allopolyploid genome of *M. javanica*

We have used long-read sequencing and modern bioinformatic approaches to assemble and phase the allopolyploid genome of the plant pathogenic nematode *Meloidogyne javanica*. Our goal was to assemble a contiguous diploid assembly for *M. javanica*, representing the A and B subgenomes separately. Our current diploid assembly (150,545,692 bp) is, as expected for a tetraploid, approximately half the 297 ±27Mb measured by flow cytometry <u>[19,28]</u>. This total length is representative of both A and B subgenomes except for collapsed regions where sequence for both subgenomes A and B is considered together. Because some regions of the assembly have been shown by k-mer profiling and coverage analysis to be present in less than four copies (Fig 1; S8 Fig), splitting the subgenomes into their component copies (A1, A2, B1, B2) to create a tetraploid assembly would not be expected to completely double the length.

#### Annotation

We annotated our assembly using both *ab initio* feature prediction algorithms and mapping of the full-length transcript sequences (S3 Table). The total number of genes predicted - 22,433 - is within the range expected for a MIG species and is comparable to previous *M. javanica* assemblies <u>[19,28]</u>.

BUSCO scores for *Meloidogyne* are consistently lower than those of more widely studied organisms, and the number of genes we have detected (S3 Table) in the genome is consistent with what has been found for other *Meloidogyne* species <u>[19,28,46]</u>. The paucity of established protein databases for less frequently investigated genera limits the accuracy of prediction-based annotation methods. Loss of core genes are not unexpected for this group of highly specialised, obligate parasites. With increased availability of sequence resources for plant parasitic nematodes, it should soon be possible to develop a more appropriate set of core genes.

#### Evidence for a chromosome-scale assembly

Previous genomic assemblies of *M. javanica* have contig counts numbering in the thousands <u>[19,28]</u>. In our current assembly, more than 99% of reads in our PacBio HiFi libraries map to 33 large scaffolds (Fig 2B). We propose that most of the 33 scaffolds represent full-length or nearly full-length chromosomes. Cytological examination in *M. javanica* indicates that the chromosome number ranges from 42-48 due to variation between isolates (Eisenback and Triantaphyllou, 1991; Triantaphyllou 1985, Janssen et al 2017). Thus, we would expect 21-24 scaffolds in our assembly. The discrepancy between scaffold number and cytological observations could be due to an imperfect assembly or failure to identify very small chromosomes in the cytological studies. We have employed several independent scaffolding softwares, Hi-C chromatin contact mapping, and the manual examination of long-read mapping to contig termini, and see no evidence to support fusing additional contigs. Additional molecular and cytological studies may be required to resolve these differences.

Many chromosome-scale assemblies identify telomeres as defining the range of their scaffolds, yet we did not identify canonical telomeric repeats at scaffold termini. Additionally, we were also not able to identify a homolog of *C. elegans* telomerase (*trt-1;* ACC: NM_001373211.4) in our assembly. This may suggest that non-standard telomere processes might be operating in root-knot nematodes as has been found for fruit flies <u>[49,50]</u>.

### Genetic variation in *M. javanica* is dominated by indels

We present the assembly as a diploid representation. Read depth analysis of individual scaffolds indicates that homologs are not identical and indels are frequent. Additionally some scaffolds appear to be present as a single copy suggesting that one of the homologs may have been lost. We identified many indels that delete or disrupt one or more copies of coding sequences, indicating that *M. javanica* is no longer genome-wide tetraploid either in copy number or functionality. A higher propensity for indel accumulation has been frequently seen in hybrid parthenogenetic species <u>[51]</u> and partial return to lower ploidy is a characteristic of many polyploids.

We find that 87% of our diploid assembly consists of one copy of either subgenome in homoeologous pairs, with two copies mapping to phased scaffolds and four copies mapping to collapsed regions. We successfully phased much of our assembly into subgenomes using k-mer signatures, enabling for the first time initial genome-wide comparison of MIG A and B subgenomes. Some scaffolds, notably scaffold 2 and some of the short scaffolds, could not be assigned to subgenomes. Scaffold 2 displays four levels of stratification in its coverage (S8 Fig) and four peaks in its depth distribution (Fig 3b) indicating that the four copies (A1+A2 and B1+B2) are almost entirely collapsed into a single scaffold. We suggest that the homoeologous chromosomes represented by scaffold 2 may be too similar in sequence to assign to a subgenome by k-mer analysis due to a possible homogenization event. For the smaller scaffolds, the failure may have been due to their small size because they did not contain enough relevant k-mers. Some of the small scaffolds were later assigned to a subgenome using transcript alignments. No misassembly or erroneous scaffolding of the unphased sequences was detected through either programmatic or manual methods.

The allotetraploid genome of *M. javanica* demonstrates extensive synteny between the A and B subgenomes with most genic regions represented in four copies. We would expect however that there would be differences between the subgenomes, including indels and other structural variation, as this is typically observed between different species and the MIG have a hybrid origin. In accordance with this, we observed very substantial structural variation between subgenomes A and B, including insertions, deletions, and translocations (Fig 4).

### Loss of synteny and fragmentation

We observe regions of the *M. javanica* genome where synteny between paired chromosomes is disrupted. One reason for this could be that the parental species (A and B) had genomes in which the non-syntenic regions had diverged by translocation, insertion or deletion. Upon hybridisation to create the allopolyploid MIG these diverged regions form the end of synteny blocks. An alternative explanation is that these changes happened after the hybridisation event in the tumultuous process of genome stabilisation immediately following it <u>[52,53]</u>. Hybridisation, polyploidization, and the loss of meiosis are processes often associated with rapid genomic change <u>[54][51,54,55]</u> and the unique MIG species allopolyploid genomes we are currently studying may represent different balances between these forces. We observe 11 chromosome-scale scaffolds in subgenome A and 17 in subgenome B despite the clear synteny throughout these two genomes. Several scaffolds in subgenome A contain blocks of genes with regions that are syntenic to different subgenome B scaffolds. Similarly, there are cases where syntenic blocks in subgenome B are present on different scaffolds in subgenome A. Together these differences suggest that ancestral chromosomal fission, or fusion events or other types of exchange have occurred. *Meloidogyne* species, like other nematodes, have holocentric chromosomes (Triantaphyllou, 1985). Genomes with dispersed centromere structure are predicted to better tolerate chromosome fragmentation and fusion <u>[56]</u>, as are species with ameiotic mechanisms of reproduction. The observed differences in copy number of chromosomes between isolates of *M. javanica* may be additional evidence for tolerance of chromosome fragmentation/fusion. Although it is always necessary to consider the possibility of misassembly or rogue scaffolding, we have substantial evidence through manual inspection and contiguous mapping of ultra-long Oxford Nanopore reads that our assembly is correct.

The majority of published assemblies of allopolyploids come from plants, where polyploidy might have shaped the genomes of around 70% of species <u>[57]</u>. Many allopolyploid plant genomes, however, show a higher level of synteny and structural conservation than we observe for *M. javanica* <u>[13,45,58]</u>. Similarly the few available chromosomal allopolyploid animal genome sequences available <u>[14,15]</u> do not show extensive deletions and chromosomal fissions as does our genome. Unlike other allopolyploid species with chromosomal genome sequences, *Meloidogyne javanica* reproduces by obligatory mitotic parthenogenesis (apomixis) and this lack of meiotic chromosome pairing may allow greater structural divergence and tolerance for the decay of synteny. We note that genomes from species not able to reproduce by meiosis are currently rare <u>[51]</u> and suggest that much more substantial genomic work on a range of species with different reproductive modes and ploidy levels will be required to reveal the diverse mechanisms shaping these genomes. It is apparent however that the changes surrounding allopolyploidy have contributed to the gene content, heterozygosity, and copy number throughout the *M. javanica* genome and these processes of locally fixed heterozygosity or rediploidization may contribute extensively to adaptive functional variation <u>[59]</u>.

### Regional homogenization of subgenomes

Some regions of the genome do not phase into A and B homoeologs due to very low divergence between the four gene copies. This could be explained either by the loss of whole chromosomes, compensated by the duplication of the remaining chromosome, or mitotic recombination (gene conversion) between homoeologs <u>[60,61]</u>. It is unclear from this single genome how asexual recombination contributes to shaping the diversity of *M. javanica*, however this has been suggested in previous MIG genomic studies <u>[19,28]</u> and may be further elucidated by our ongoing molecular evolution and population genomic studies.

The initial tetraploidy of *M. javanica* will buffer against deleterious phenotypic consequences of indels, allowing a substantial accumulation of genetic variation by this mechanism. It has been argued for angiosperms that this rediploidization process can contribute to adaptive species divergence by providing genomic and transcriptomic diversity <u>[59,62]</u>. Other mechanisms of adaptive divergence may also operate at the same time. The increase in gene copy number created by polyploidization gives the potential for functional gene divergence by neo- or sub-functionalization, as well as adaptive phenotypes driven by copy number loss <u>[63–65]</u>. Adaptation by gene copy number variation has already been reported in *M. incognita* <u>[66]</u> and it may be that genomic copy number variation more broadly is a major source of functional genetic variation in the MIG.

### A genomic framework for RKN functional and diversity studies

In this paper we present a highly contiguous, annotated and phased genome of the allotetraploid plant pathogenic nematode *M. javanica*. This genome assembly will provide many tools for diverse investigations by plant pathologists and nematologists in addition to adding to our understanding of the origins and diversity of *M. javanica*. It will also serve as a reference for investigating genome structure and pathogenicity in other *Meloidogyne* species.

The contiguous nature of this genome and the high quality annotation will facilitate RKN functional studies since transcripts can be mapped accurately to the annotated subgenome. This allows consideration of copy number variation, which may be an important component of functional variation in these species.

Progress is being made by many groups in understanding the basis of nematode virulence and the key loci involved <u>[67–69]</u>. In such cases even light coverage sequencing of field isolates mapped to a high quality genome could give valuable information to commercial growers about the likely pathogenicity of those strains <u>[70]</u>.

## Methods

### Reproducibility

Wherever possible, this study attempted to contain all bioinformatic processes in reproducible workflows or scripts, for the purpose of openness and enabling replication. Workflows and code are archived in a Zenodo repository along with final outputs (doi://XXXXXX.XX). The raw reads plus final nuclear and mitochondrial assemblies are available on the International Nucleotide Sequence Databases (INSDC) (ACC: XXXXXX) (ACC: XXXXXX).

### Biological material

*Meloidogyne javanica* strain VW4 was used for this work (Gleason et al, 2008; Szitenberg et al., 2017). Cultures of this strain have been maintained on tomato plants under greenhouse conditions for over 30 years (Yaghoobi et al, 1995). Periodic transfers of single egg masses have been carried out to maintain uniformity. For DNA preparation, eggs were harvested from roots and cleaned by sucrose flotation as previously described (Branch et al, 2004) then flash-frozen in liquid N2. High molecular weight DNA (HMW DNA) isolation was carried out at UC Davis Genome Center (Supplementary Methods). Integrity of the HMW gDNA was verified on a Femto Pulse system (Agilent Technologies, CA) where majority of the DNA was found to be in fragments above 100 Kb.

Total RNA was isolated from three *M. javanica* life stages: eggs, freshly hatched juveniles, and females dissected from tomato roots 21 days after infection. This material was flash-frozen and RNA was extracted using an RNeasy Kit (Qiagen, USA) following the manufacturer’s instructions. TURBO DNase treatment was carried out to remove genomic DNA from total RNA samples (TURBO DNA-free Kit™, Ambion, USA). RNA concentration and purity were measured using a NanoDrop OneC Microvolume UV-Vis Spectrophotometer (Thermo Scientific, USA). RNA integrity and quality was assessed using bioanalyzer.

### Sequencing and QC

#### High fidelity (HiFi) long-read sequencing

PacBio HiFi library preparation and sequencing of HMW DNA was performed by UC Davis DNA Technologies Core on a PacBio Sequel II (Supplementary Methods). Data from the two generated libraries were pooled and only reads longer than 5000 bp and with a quality score over 15 were retained.

#### Nanopore sequencing

Nanopore sequencing of HMW DNA using Oxford Nanopore Technologies (ONT) systems was carried out by UC Davis DNA Technologies Core. The super-long-read DNA sequencing protocol (Supplementary Methods) yielded 23 gbp of data, which was then filtered to contain only reads longer than 25 kbp.

#### Hi-C chromatin conformation capture

DNA was prepared using Proximo Hi-C kit (Animal) as recommended by the manufacturer (Phase Genomics, Seattle, WA, USA). Library preparation and sequencing were carried out at the UC Davis DNA Technologies Core and scaffolding with Proximo was performed by Phase Genomics (Supplementary Methods).

#### PacBio Iso-Seq

Reads from one Sequel II SMRT cell were quality controlled and converted into clustered reads using the *IsoSeq3* pipeline with default parameters <u>[71]</u>.

### Genome profiling

Profiling was performed by *Genomescope2* and *smudgeplot <u>[38]</u>.* All quality controlled PacBio HiFi libraries were combined and used as input for these programs. Both were run with default parameters, aside from -*ploidy* in *Genomescope2* set to 4, as this was the primary ploidy indicated by *smudgeplot* and suggested by visual analysis of mapped reads.

### Assembly

#### Draft assembly

Initial draft assemblies were generated using several different assemblers and appraised with *asmapp <u>[72]</u>.* The most appropriate assembly based on length and contiguity was carried forward, after which haplotigs were identified and removed using *purge_dups* <u>[73]</u>.

#### Scaffolding with Oxford Nanopore and Hi-C

We used *SLR* <u>[74]</u> to scaffold the assembly with our trimmed and concatenated ONT reads. The assembly was then scaffolded with Hi-C reads using the *Proximo* pipeline by Phase Genomics Ltd. The resulting contact map was manually curated using *Juicebox* assembly tools <u>[75]</u> to produce the most likely assembly based on contact linkage information and TAD presence or absence. A third phase of scaffolding was performed by *samba,* to break potential misassemblies introduced during manual *Juicebox* scaffolding and scaffold sequences that were broken as debris <u>[40]</u>.

### Annotation

#### Repeat annotation

*RepeatModeler* was applied to the assembly <u>[76]</u>, with the resulting library of repeat models used as input for *RepeatMasker* to annotate repeat regions and generate a soft-masked version of the genome <u>[77]</u>.

#### Gene annotation

Iso-Seq reads were mapped to the genome and collapsed using a *snakemake* workflow automating the *IsoSeq3* pipeline <u>[71]</u>. The annotation pipeline *MAKER3* was then employed to perform *ab initio* and predictive annotation of our genome <u>[78,79]</u>. Full methods of annotation iterations and parameters can be found in Supplementary Methods.

### Subgenome phasing

#### Identification of homoeologous scaffold pairs

Homoeologous pairs were detected through identification of orthologs shared between scaffolds (Supplementary Methods). Pairs sharing a large number of orthologs and nucleotide similarity were considered homoeologous. This was validated with other methods, including MASH distance <u>[80]</u> and shared possession of duplicated BUSCO genes <u>[81]</u>.

#### Phasing of subgenomes

Scaffolds were phased into A and B subgenomes using a k-mer-based approach built on the approach taken by <u>[45]</u>. K-mers in the assembly were detected and counted using *jellyfish<u>[82]</u>*, with only k-mers present more than 75 times in the assembly and represented at least twice as often in one subgenome than the other carried forward. These counts were then transformed into binomial distributions. Hierarchical clustering was performed on these sets, creating a dendrogram placing scaffolds into opposing clusters, each cluster representing a subgenome (Supplementary Methods).

### Synteny analysis

The *MCScan (Python)* <u>[83]</u> module of *JCVI <u>[84]</u>*was used to perform a synteny analysis between the two subgenomes. Custom scripts then extracted collinearity information and generated synteny plots (Supplementary Methods).

## Acknowledgments

We acknowledge the Viper High Performance Computing facility of the University of Hull and its support team, specifically Chris Collins, for their invaluable help in managing and deploying software, as well as the high Performance Computing team at UC Davis for making available their resources for part of our analysis.

## Supporting information

**S1 Fig: Length and quality statistics of concatenated and quality controlled PacBio HiFi libraries.**

**S2 Fig: Length and quality statistics of concatenated and quality controlled Oxford Nanopore libraries.**

**S3 Fig: Total cumulative length over number of scaffolds.**

**S4 Fig: Blobplot of contamination in our *Meloidogyne javanica* assembly**

**S5 Fig: BUSCO v5 results.**

**S6 Fig: Assembly-wide distributions of coverage**

**S7 Fig: Scaffold-level distributions of coverage**

**S8 Fig: Coverage depth of individual bases across scaffolds.**

**S9 Fig: Subgenome phased dendrogram of scaffolds.**

**S1 Table: Descriptive statistics of assembly and its contemporaries**

**S2 Table: Tabular breakdown of repeat annotation from *RepeatMasker*.**

**S3 Table: Full table of results from structural annotation with MAKER3.**

**S4 Table: List of pairs that share CDS and amount of shared links.**

**S5 Table: Validation of homoeologous pairings.**

**S6 Table: Summary of the longest 33 scaffolds.**

## References

1. Glover NM, Redestig H, Dessimoz C. Homoeologs: What Are They and How Do We Infer Them? Trends Plant Sci. 2016;21: 609–621. doi:10.1016/j.tplants.2016.02.005

2. Barker MS, Arrigo N, Baniaga AE, Li Z, Levin DA. On the relative abundance of autopolyploids and allopolyploids. New Phytol. 2015. doi:10.1111/nph.13698

3. Schoenfelder KP, Fox DT. The expanding implications of polyploidy. J Cell Biol. 2015;209: 485–491. doi:10.1083/jcb.201502016

4. Session AM, Uno Y, Kwon T, Chapman JA, Toyoda A, Takahashi S, et al. Genome evolution in the allotetraploid frog Xenopus laevis. Nature. 2016;538: 336–343. doi:10.1038/nature19840

5. Ming R, Man Wai C. Assembling allopolyploid genomes: no longer formidable. Genome Biol. 2015;16: 27. doi:10.1186/s13059-015-0585-5

6. Kyriakidou M, Tai HH, Anglin NL, Ellis D, Strömvik MV. Current Strategies of Polyploid Plant Genome Sequence Assembly. Front Plant Sci. 2018;9: 1660. doi:10.3389/fpls.2018.01660

7. Rhie A, McCarthy SA, Fedrigo O, Damas J, Formenti G, Koren S, et al. Towards complete and error-free genome assemblies of all vertebrate species. Nature. 2021;592: 737–746. doi:10.1038/s41586-021-03451-0

8. Zhang X, Wu R, Wang Y, Yu J, Tang H. Unzipping haplotypes in diploid and polyploid genomes. Comput Struct Biotechnol J. 2020;18: 66–72. doi:10.1016/j.csbj.2019.11.011

9. Saada OA, Friedrich A, Schacherer J. Towards accurate, contiguous and complete alignment-based polyploid phasing algorithms. Genomics. 2022;114: 110369. doi:10.1016/j.ygeno.2022.110369

10. Edger PP, Poorten TJ, VanBuren R, Hardigan MA, Colle M, McKain MR, et al. Origin and evolution of the octoploid strawberry genome. Nat Genet. 2019;51: 541–547. doi:10.1038/s41588-019-0356-4

11. Gan X, Li S, Zong Y, Cao D, Li Y, Liu R, et al. Chromosome-Level Genome Assembly Provides New Insights into Genome Evolution and Tuberous Root Formation of Potentilla anserina. Genes. 2021;12. doi:10.3390/genes12121993

12. Kolesnikova UK, Scott AD, Van de Velde JD, Burns R, Tikhomirov NP, Pfordt U, et al. Genome of selfing Siberian Arabidopsis lyrata explains establishment of allopolyploid Arabidopsis kamchatica. bioRxiv. 2022. p. 2022.06.24.497443. doi:10.1101/2022.06.24.497443

13. Zheng Y, Yang D, Rong J, Chen L, Zhu Q, He T, et al. Allele-aware chromosome-scale assembly of the allopolyploid genome of hexaploid Ma bamboo (Dendrocalamus latiflorus Munro). J Integr Plant Biol. 2022;64: 649–670. doi:10.1111/jipb.13217

14. Du K, Stöck M, Kneitz S, Klopp C, Woltering JM, Adolfi MC, et al. The sterlet sturgeon genome sequence and the mechanisms of segmental rediploidization. Nature Ecology & Evolution. 2020;4: 841–852. doi:10.1038/s41559-020-1166-x

15. Kuhl H, Du K, Schartl M, Kalous L, Stöck M, Lamatsch DK. Equilibrated evolution of the mixed auto-/allopolyploid haplotype-resolved genome of the invasive hexaploid Prussian carp. Nat Commun. 2022;13: 4092. doi:10.1038/s41467-022-31515-w

16. Perry RN, Moens M, Starr JL. Root-knot Nematodes. CABI; 2009. Available: https://play.google.com/store/books/details?id=UN3uHMr_UCoC

17. Williamson VM, Gleason CA. Plant–nematode interactions. Curr Opin Plant Biol. 2003;6: 327–333. doi:10.1016/S1369-5266(03)00059-1

18. Trudgill DL, Blok VC. Apomictic, polyphagous root-knot nematodes: exceptionally successful and damaging biotrophic root pathogens. Annu Rev Phytopathol. 2001;39: 53–77. doi:10.1146/annurev.phyto.39.1.53

19. Szitenberg A, Salazar-Jaramillo L, Blok VC, Laetsch DR, Joseph S, Williamson VM, et al. Comparative Genomics of Apomictic Root-Knot Nematodes: Hybridization, Ploidy, and Dynamic Genome Change. Genome Biol Evol. 2017;9: 2844–2861. doi:10.1093/gbe/evx201

20. Wesemael W, Viaene N, Moens M. Root-knot nematodes (Meloidogyne spp.) in Europe. Nematology. 2011;13: 3–16. doi:10.1163/138855410X526831

21. Jones JT, Haegeman A, Danchin EGJ, Gaur HS, Helder J, Jones MGK, et al. Top 10 plant-parasitic nematodes in molecular plant pathology. Mol Plant Pathol. 2013;14: 946–961. Available: https://bsppjournals.onlinelibrary.wiley.com/doi/abs/10.1111/mpp.12057

22. Bernard GC, Egnin M, Bonsi C. The impact of plant-parasitic nematodes on agriculture and methods of control. Nematology-Concepts, Diagnosis and Control. 2017;10. Available: https://books.google.co.uk/bookshl=en&lr=&id=MQWQDwAAQBAJ&oi=fnd&pg=PA121&dq=agricultural+impact+root-knot+nematodes&ots=bBkbCxpflc&sig=1tg90hY-8Y2Z7V_6F8KGljCENoM

23. Eisenback JD, Triantaphyllou HH. Root-knot nematodes: Meloidogyne species and races. Manual of agricultural nematology. 1991;1: 191–274. Available: https://www.researchgate.net/profile/Jonathan_Eisenback/publication/283548298_Root-Knot_Nematodes_Meloidogyne_Species_and_Races/links/563e67e308ae34e98c4d93c1/Root-Knot-Nematodes-Meloidogyne-Species-and-Races.pdf

24. Subbotin SA, Rius JEP, Castillo P. Systematics of Root-knot Nematodes (Nematoda: Meloidogynidae). BRILL; 2021. Available: https://play.google.com/store/books/details?id=xs87EAAAQBAJ

25. Triantaphyllou AC. Gametogenesis and the Chromosomes of Meloidogyne nataliei: Not Typical of Other Root-knot Nematodes. J Nematol. 1985;17: 1–5. Available: https://www.ncbi.nlm.nih.gov/pubmed/19294050

26. Bird DM, Williamson VM, Abad P, McCarter J, Danchin EGJ, Castagnone-Sereno P, et al. The genomes of root-knot nematodes. Annu Rev Phytopathol. 2009;47: 333–351. doi:10.1146/annurev-phyto-080508-081839

27. Lunt DH. Genetic tests of ancient asexuality in root knot nematodes reveal recent hybrid origins. BMC Evol Biol. 2008;8: 194. doi:10.1186/1471-2148-8-194

28. Blanc-Mathieu R, Perfus-Barbeoch L, Aury J-M, Da Rocha M, Gouzy J, Sallet E, et al. Hybridization and polyploidy enable genomic plasticity without sex in the most devastating plant-parasitic nematodes. PLoS Genet. 2017;13: e1006777. doi:10.1371/journal.pgen.1006777

29. Hartman KM, Sasser JN. Identification of Meloidogyne species on the basis of differential host test and perineal pattern morphology. An advanced treatise on Meloidogyne. 1985;2: 69–77. Available: https://books.google.com/books?hl=en&lr=&id=gNo4AQAAIAAJ&oi=fnd&pg=PA69&dq=hartman+and+sasser&ots=r0T3kOF2TR&sig=qp5-qepxxIy-MaZ3Yxq8q3OTlBg

30. Roberts PA, Thomason J. Variability in reproduction of isolates of Meloidogyne incognita and M. javanica on resistant tomato genotypes. worldveg.tind.io; 1986. Available: https://worldveg.tind.io/record/4772/

31. Rammah A, Hirschmann H. Morphological Comparison of Three Host Races of Meloidogyne javanica. J Nematol. 1990;22: 56–68. Available: https://www.ncbi.nlm.nih.gov/pubmed/19287689

32. Williamson VM, Kumar A. Nematode resistance in plants: the battle underground. Trends Genet. 2006;22: 396–403. doi:10.1016/j.tig.2006.05.003

33. Gleason CA, Liu QL, Williamson VM. Silencing a candidate nematode effector gene corresponding to the tomato resistance gene Mi-1 leads to acquisition of virulence. Mol Plant Microbe Interact. 2008;21: 576–585. doi:10.1094/MPMI-21-5-0576

34. Hajihassani A, Marquez J, Woldemeskel M, Hamidi N. Identification of Four Populations of Meloidogyne incognita in Georgia, United States, Capable of Parasitizing Tomato-Bearing Mi-1.2 Gene. Plant Dis. 2022;106: 137–143. doi:10.1094/PDIS-05-21-0902-RE

35. Sato K, Kadota Y, Gan P, Bino T, Uehara T, Yamaguchi K, et al. High-Quality Genome Sequence of the Root-Knot Nematode Meloidogyne arenaria Genotype A2-O. Genome Announc. 2018;6. doi:10.1128/genomeA.00519-18

36. Susič N, Koutsovoulos GD, Riccio C, Danchin EGJ, Blaxter ML, Lunt DH, et al. Genome sequence of the root-knot nematode Meloidogyne luci. J Nematol. 2020;52: 1–5. doi:10.21307/jofnem-2020-025

37. Fox DT, Soltis DE, Soltis PS, Ashman T-L, Van de Peer Y. Polyploidy: A biological force from cells to ecosystems. Trends Cell Biol. 2020;30: 688–694. doi:10.1016/j.tcb.2020.06.006

38. Ranallo-Benavidez TR, Jaron KS, Schatz MC. GenomeScope 2.0 and Smudgeplot for reference-free profiling of polyploid genomes. Nat Commun. 2020;11: 1432. doi:10.1038/s41467-020-14998-3

39. Cheng H, Concepcion GT, Feng X, Zhang H, Li H. Haplotype-resolved de novo assembly using phased assembly graphs with hifiasm. Nat Methods. 2021;18: 170–175. doi:10.1038/s41592-020-01056-5

40. Zimin AV, Salzberg SL. The SAMBA tool uses long reads to improve the contiguity of genome assemblies. PLoS Comput Biol. 2022;18: e1009860. doi:10.1371/journal.pcbi.1009860

41. Laetsch DR, Blaxter ML. BlobTools: Interrogation of genome assemblies. F1000Res. 2017;6: 1287. doi:10.12688/f1000research.12232.1

42. Parra G, Bradnam K, Korf I. CEGMA: a pipeline to accurately annotate core genes in eukaryotic genomes. Bioinformatics. 2007;23: 1061–1067. doi:10.1093/bioinformatics/btm071

43. Koutsovoulos GD, Poullet M, El Ashry A, Kozlowski DK, Sallet E, Da Rocha M, et al. The polyploid genome of the mitotic parthenogenetic root-knot nematode Meloidogyne enterolobii. bioRxiv. 2019. p. 586818. doi:10.1101/586818

44. Kozlowski DKL, Hassanaly-Goulamhoussen R, Da Rocha M, Koutsovoulos GD, Bailly-Bechet M, Danchin EGJ. Movements of transposable elements contribute to the genomic plasticity and species diversification in an asexually reproducing nematode pest. Evol Appl. 2021;14: 1844–1866. doi:10.1111/eva.13246

45. Cerca J, Petersen B, Lazaro-Guevara JM, Rivera-Colón A, Birkeland S, Vizueta J, et al. The genomic basis of the plant island syndrome in Darwin’s giant daisies. Nat Commun. 2022;13: 3729. doi:10.1038/s41467-022-31280-w

46. Koutsovoulos GD, Poullet M, Elashry A, Kozlowski DKL, Sallet E, Da Rocha M, et al. Genome assembly and annotation of Meloidogyne enterolobii, an emerging parthenogenetic root-knot nematode. Scientific Data. 2020;7: 324. doi:10.1038/s41597-020-00666-0

47. Koutsovoulos GD, Marques E, Arguel M, Duret L, Machado ACZ, Carneiro RMDG, et al. Population genomics supports clonal reproduction and multiple independent gains and losses of parasitic abilities in the most devastating nematode pest. Evol Appl. 2019;26: 909. doi:10.1111/eva.12881

48. Bali S, Hu S, Vining K, Brown C, Mojtahedi H, Zhang L, et al. Nematode genome announcement: Draft genome of Meloidogyne chitwoodi, an economically important pest of potato in the pacific northwest. Mol Plant Microbe Interact. 2021; MPMI12200337A. doi:10.1094/MPMI-12-20-0337-A

49. Mason JM, Reddy HM, Frydrychova RC. Telomere Maintenance in Organisms without Telomerase. In: Seligmann H, editor. DNA Replication. Rijeka: IntechOpen; 2011. doi:10.5772/19348

50. Pardue M-L, DeBaryshe PG. Retrotransposons that maintain chromosome ends. Proc Natl Acad Sci U S A. 2011;108: 20317–20324. doi:10.1073/pnas.1100278108

51. Jaron KS, Bast J, Nowell RW, Ranallo-Benavidez TR, Robinson-Rechavi M, Schwander T. Genomic Features of Parthenogenetic Animals. J Hered. 2021;112: 19–33. doi:10.1093/jhered/esaa031

52. Edger PP, McKain MR, Bird KA, VanBuren R. Subgenome assignment in allopolyploids: challenges and future directions. Curr Opin Plant Biol. 2018;42: 76–80. doi:10.1016/j.pbi.2018.03.006

53. Emery M, Willis MMS, Hao Y, Barry K, Oakgrove K, Peng Y, et al. Preferential retention of genes from one parental genome after polyploidy illustrates the nature and scope of the genomic conflicts induced by hybridization. PLoS Genet. 2018;14: e1007267. doi:10.1371/journal.pgen.1007267

54. Ma X-F, Gustafson JP. Genome evolution of allopolyploids: a process of cytological and genetic diploidization. Cytogenet Genome Res. 2005;109: 236–249. doi:10.1159/000082406

55. Balloux F, Lehmann L, de Meeûs T. The Population Genetics of Clonal and Partially Clonal Diploids. Genetics. 2003;164: 1635–1644. doi:10.1093/genetics/164.4.1635

56. Carlton PM, Davis RE, Ahmed S. Nematode chromosomes. Genetics. 2022. doi:10.1093/genetics/iyac014

57. Masterson J. Stomatal size in fossil plants: evidence for polyploidy in majority of angiosperms. Science. 1994;264: 421–424. doi:10.1126/science.264.5157.421

58. Shen Y, Li W, Zeng Y, Li Z, Chen Y, Zhang J, et al. Chromosome-level and haplotype-resolved genome provides insight into the tetraploid hybrid origin of patchouli. Nat Commun. 2022;13: 1–15. doi:10.1038/s41467-022-31121-w

59. Dodsworth S, Chase MW, Leitch AR. Is post-polyploidization diploidization the key to the evolutionary success of angiosperms? Bot J Linn Soc. 2016;180: 1–5. doi:10.1111/boj.12357

60. Harris S, Rudnicki KS, Haber JE. Gene conversions and crossing over during homologous and homeologous ectopic recombination in Saccharomyces cerevisiae. Genetics. 1993;135: 5–16. doi:10.1093/genetics/135.1.5

61. Mansai SP, Innan H. The power of the methods for detecting interlocus gene conversion. Genetics. 2010;184: 517–527. doi:10.1534/genetics.109.111161

62. Hollister JD. Polyploidy: adaptation to the genomic environment. New Phytol. 2015;205: 1034–1039. doi:10.1111/nph.12939

63. Paquin C, Adams J. Frequency of fixation of adaptive mutations is higher in evolving diploid than haploid yeast populations. Nature. 1983;302: 495–500. doi:10.1038/302495a0

64. Otto SP, Whitton J. Polyploid incidence and evolution. Annu Rev Genet. 2000;34: 401–437. doi:10.1146/annurev.genet.34.1.401

65. Zörgö E, Chwialkowska K, Gjuvsland AB, Garré E, Sunnerhagen P, Liti G, et al. Ancient evolutionary trade-offs between yeast ploidy states. PLoS Genet. 2013;9: e1003388. doi:10.1371/journal.pgen.1003388

66. Castagnone-Sereno P, Mulet K, Danchin EGJ, Koutsovoulos GD, Karaulic M, Da Rocha M, et al. Gene copy number variations as signatures of adaptive evolution in the parthenogenetic, plant-parasitic nematode Meloidogyne incognita. Mol Ecol. 2019;26: 906. doi:10.1111/mec.15095

67. Pogorelko GV, Juvale PS, Rutter WB, Hütten M, Maier TR, Hewezi T, et al. Re-targeting of a plant defense protease by a cyst nematode effector. Plant J. 2019;98: 1000–1014. doi:10.1111/tpj.14295

68. Kihika R, Tchouassi DP, Ng’ang’a MM, Hall DR, Beck JJ, Torto B. Compounds Associated with Infection by the Root-Knot Nematode, Meloidogyne javanica, Influence the Ability of Infective Juveniles to Recognize Host Plants. J Agric Food Chem. 2020;68: 9100–9109. doi:10.1021/acs.jafc.0c03386

69. Song H, Lin B, Huang Q, Sun T, Wang W, Liao J, et al. The Meloidogyne javanica effector Mj2G02 interferes with jasmonic acid signalling to suppress cell death and promote parasitism in Arabidopsis. Mol Plant Pathol. 2021;22: 1288–1301. doi:10.1111/mpp.13111

70. Sellers GS, Jeffares DC, Lawson B, Prior T, Lunt DH. Identification of individual root-knot nematodes using low coverage long-read sequencing. PLoS One. 2021;16: e0253248. doi:10.1371/journal.pone.0253248

71. PacificBiosciences. IsoSeq3. Github; 2022. Available: https://github.com/PacificBiosciences/IsoSeq

72. Winter M. asmapp: ASMAPP assembly appraisal workflow. Built in snakemake, ASMAPP performs many basic and intermediate assembly appraisal tasks. Github; 2022. Available: https://github.com/mrmrwinter/asmapp

73. Guan D, McCarthy SA, Wood J, Howe K, Wang Y, Durbin R. Identifying and removing haplotypic duplication in primary genome assemblies. Bioinformatics. 2020;36: 2896–2898. doi:10.1093/bioinformatics/btaa025

74. Luo J, Lyu M, Chen R, Zhang X, Luo H, Yan C. SLR: a scaffolding algorithm based on long reads and contig classification. BMC Bioinformatics. 2019;20: 1–11. doi:10.1186/s12859-019-3114-9

75. Durand NC, Robinson JT, Shamim MS, Machol I, Mesirov JP, Lander ES, et al. Juicebox Provides a Visualization System for Hi-C Contact Maps with Unlimited Zoom. Cell Syst. 2016;3: 99–101. doi:10.1016/j.cels.2015.07.012

76. Smit AFA, Hubley R, Green P. RepeatModeler Open-1.0. 2008--2015. Seattle, USA: Institute for Systems Biology Available from: http://www.repeatmasker.org, Last Accessed May. 2015;1: 2018.

77. Smit AFA, Hubley R, Green P. RepeatMasker Open-4.0. 2013--2015. 2015.

78. Holt C, Yandell M. MAKER2: an annotation pipeline and genome-database management tool for second-generation genome projects. BMC Bioinformatics. 2011;12: 491. doi:10.1186/1471-2105-12-491

79. Campbell MS, Holt C, Moore B, Yandell M. Genome Annotation and Curation Using MAKER and MAKER-P. Curr Protoc Bioinformatics. 2014;48: 4.11.1–39. doi:10.1002/0471250953.bi0411s48

80. Ondov BD, Treangen TJ, Melsted P, Mallonee AB, Bergman NH, Koren S, et al. Mash: fast genome and metagenome distance estimation using MinHash. Genome Biol. 2016;17: 132. doi:10.1186/s13059-016-0997-x

81. Simão FA, Waterhouse RM, Ioannidis P, Kriventseva EV, Zdobnov EM. BUSCO: assessing genome assembly and annotation completeness with single-copy orthologs. Bioinformatics. 2015;31: 3210–3212. doi:10.1093/bioinformatics/btv351

82. Marçais G, Kingsford C. A fast, lock-free approach for efficient parallel counting of occurrences of k-mers. Bioinformatics. 2011;27: 764–770. doi:10.1093/bioinformatics/btr011

83. Tang H, Bowers JE, Wang X, Ming R, Alam M, Paterson AH. Synteny and collinearity in plant genomes. Science. 2008;320: 486–488. doi:10.1126/science.1153917

84. Tang H, Krishnakumar V, Li J. jcvi: JCVI utility libraries. 2015. doi:10.5281/zenodo.31631

85. Altschul SF, Gish W, Miller W, Myers EW, Lipman DJ. Basic local alignment search tool. J Mol Biol. 1990;215: 403–410. doi:10.1016/S0022-2836(05)80360-2

86. Bernt M, Donath A, Jühling F, Externbrink F, Florentz C, Fritzsch G, et al. MITOS: improved de novo metazoan mitochondrial genome annotation. Mol Phylogenet Evol. 2013;69: 313–319. doi:10.1016/j.ympev.2012.08.023

87. Li H, Handsaker B, Wysoker A, Fennell T, Ruan J, Homer N, et al. The Sequence Alignment/Map format and SAMtools. Bioinformatics. 2009;25: 2078–2079. doi:10.1093/bioinformatics/btp352

88. Gurevich A, Saveliev V, Vyahhi N, Tesler G. QUAST: quality assessment tool for genome assemblies. Bioinformatics. 2013;29: 1072–1075. doi:10.1093/bioinformatics/btt086

