## Supplementary material for "Phased chromosome-scale genome assembly of an asexual, allopolyploid root-knot nematode reveals complex subgenomic structure: Analysis of the allopolyploid genome of *Meloidogyne javanica*": S1 Table.docx

#### Supplementary Table 1: Descriptive statistics of assembly and its contemporaries.

| **Accession** | **Species** | **Year** | **Scaffolds** | **Assembly size (mbp)** | **Nuclear DNA Content (mbp)** | **N50 (kbp)** | **CEGMA % (complete: C, partial: P)** | **BUSCO v3 % (complete: C, fragment: F, missing: M)** | **GC %** |
| --- | --- | --- | --- | --- | --- | --- | --- | --- | --- |
| XXXXXXXX | *M. javanica** (Hull) | 2023 | 69 | 150.5 | NA | 5,793 | C :93.95 (1.88)  P: 95.56 | C:83.9%F:6.6%; M:9.5% | 30.1 |
| GCA_003693625.1 | *M. javanica* (Hull) | 2017 | 34,394 | 142.6 | NA | 14.1 | C :89.52 (2.71)  P: 95.16 | C:87.5%F:4.3%; M:8.2% | 30.2 |
| GCA_900003945.1 | *M. javanica* (Avignon) | 2017 | 31,341 | 235.8 | 297+- 27 | 10.4 | C: 92.74 (3.68)  P :95.56 | C:90.1%  F:2.3%; M:7.6% | 30 |
| GCA_014132215.1 | *M. incognita* (Morelos) | 2017 | 12,091 | 183.5 | 189 +- 15 | 38.6 | C: 94.76 (2.93)  P :96.77 | C:88.5%  F:3.0%; M:8.5% | 29.8 |
| GCA_003693645.1 | *M. incognita* (Hull) | 2017 | 33,735 | 122 | NA | 16.5 | C: 82.66 (2.34)  P :89.52 | C:80.2%  F:7.9%; M:11.9% | 30.6 |
| GCA_000172435.1 | *M. hapla* (VW9) | 2008 | 1,523 | 53.6 | 121 +- 3 | 83.6 | C :93.55 (1.19)  P: 95.56 | C:87.4%  F:4.3%; M:8.3% | 27.4 |
| GCA_003693605.1 | *M. floridensis (SJF1)* | 2018 | 9,134 | 74.9 | NA | 13.3 | C: 77.42 (1.71)  P: 83.87 | C:76.5%  F:7.6%; M:15.9% | 30.2 |
| GCA_902706615.1 | *M. luci* (SI-Smartno) | 2020 | 327 | 209.2 | NA | 1,712 | C: 95.56 (2.92)  P :96.77 | C:87.8%  F:4.0%; M:8.2% | 30.2 |
| GCA_903994135.1 | *M. enterolobii* (Swiss) | 2021 | 4,437 | 240 | 275 +- 19 | 143 | C: 94.76 (3.30)  P: 96.77 | C:87.5%  F:3.6%; M:8.9% | 30 |
| GCA_900003985.1 | *M. arenaria (Guadeloupe)* | 2017 | 26,196 | 258.1 | 304 +- 9 | 16.5 | C :94.76 (3.66)  P :95.56 | C:87.1%  F:4.3%; M:8.6% | 30 |
| GCA_003133805.1 | *M. arenaria (A2-0)* | 2019 | 2,224 | 284.05 | NA | 204.6 | C: 94.76 (3.57)  P :96.77 | C:87.1%  F:2.6%; M:10.3% | 30 |
| GCA_002778205.2 | *M. graminicola (IARI)* | 2022 | 4,304 | 38.18 | NA | 20.4 | C: 84.27 (1.34)  P: 90.73 | C:73.6%  F:15.2%; M:11.2% | 23.05 |
