## Supplementary material for "Phased chromosome-scale genome assembly of an asexual, allopolyploid root-knot nematode reveals complex subgenomic structure: Analysis of the allopolyploid genome of *Meloidogyne javanica*": S2 Fig.docx

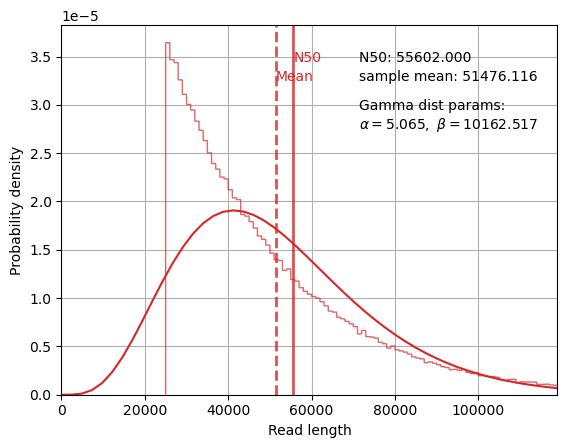

#### Supplementary Figure 2: Length and quality statistics of concatenated and quality controlled Oxford Nanopore libraries.
