## Supplementary material for "Phased chromosome-scale genome assembly of an asexual, allopolyploid root-knot nematode reveals complex subgenomic structure: Analysis of the allopolyploid genome of *Meloidogyne javanica*": S2 Table.docx

#### Supplementary Table 2: Tabular breakdown of repeat annotation from *RepeatMasker*.

|  | **Number of elements** | **Length occupies (bp)** | **Percentage of sequence (%)** |
| --- | --- | --- | --- |
| **Retroelements** | 11491 | 7443793 | 4.94 |
| SINEs | 0 | 0 | 0.00 |
| Penelope | 0 | 0 | 0.00 |
| LINEs | 663 | 657807 | 0.44 |
| - CRE/SLACS | 0 | 0 | 0.00 |
| - L2/CR1/Rex | 588 | 633246 | 0.42 |
| - R1/LOA/Jockey | 0 | 0 | 0.00 |
| - R2/R4/NeSL | 0 | 0 | 0.00 |
| - RTE/bOV-b | 0 | 0 | 0.00 |
| - L1/CIN4 | 34 | 3868 | 0.00 |
| LTR Elements | 10828 | 6785986 | 4.51 |
| - BEL/Pao | 2888 | 3074846 | 2.04 |
| - Ty1/Copia | 0 | 0 | 0.00 |
| - Gypsy/DIRS1 | 3129 | 2254025 | 1.50 |
| **DNA Transposons** | 7992 | 5674442 | 3.77 |
| Hobo-Activator | 1547 | 261731 | 0.17 |
| Tc1-IS630-Pogo | 1847 | 601027 | 0.40 |
| En-Spm | 0 | 0 | 0.00 |
| MuDR-IS905 | 0 | 0 | 0.00 |
| PiggyBac | 0 | 0 | 0.00 |
| Tourist/Harbinger | 0 | 0 | 0.00 |
| Other (Mirage, P-element, Transib) | 0 | 0 | 0.00 |
| **Rolling-circles** | 4285 | 1796873 | 1.19 |
| **Unclassified** | 129631 | 25417075 | 16.88 |
| **Total interspersed repeats** | - | 38535310 | 25.60 |
| **Small RNA** | 1092 | 1514780 | 1.01 |
| **Satellites** | 526 | 59215 | 0.04 |
| **Simple repeats** | 55163 | 2683708 | 1.78 |
| **Low complexity** | 24100 | 1270259 | 0.84 |
