## Supplementary material for "Phased chromosome-scale genome assembly of an asexual, allopolyploid root-knot nematode reveals complex subgenomic structure: Analysis of the allopolyploid genome of *Meloidogyne javanica*": S3 Fig.docx

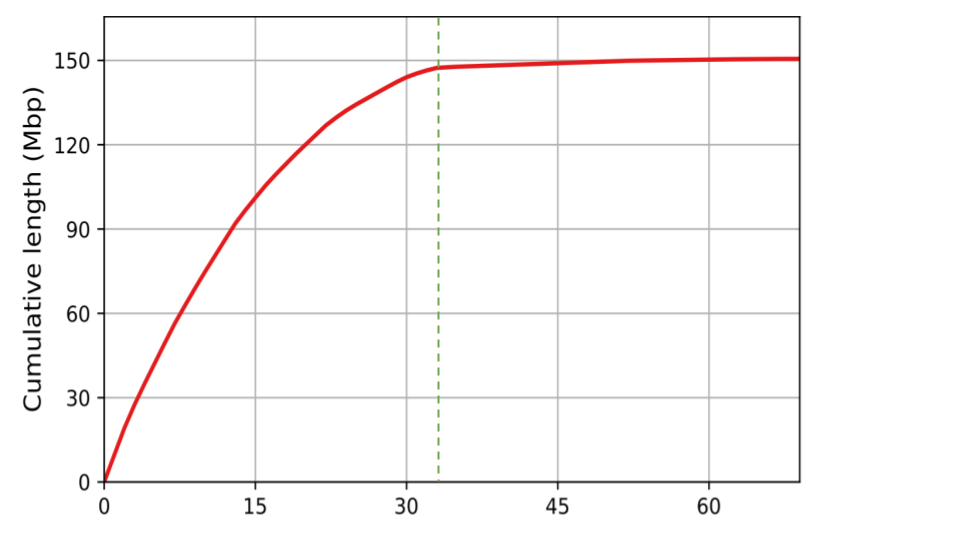

#### Supplementary Figure 3: Total cumulative length over number of scaffolds. Generated using *QUAST*. Green dashed line indicates the end of scaffold 33.
