## Supplementary material for "Phased chromosome-scale genome assembly of an asexual, allopolyploid root-knot nematode reveals complex subgenomic structure: Analysis of the allopolyploid genome of *Meloidogyne javanica*": S3 Table.docx

#### Supplementary Table 3: Full table of results from structural annotation with MAKER3.

| **Metric** | **Count** |
| --- | --- |
| Number of genes | 22433 |
| Number of mrnas | 22433 |
| Number of mrnas with utr both sides | 2811 |
| Number of mrnas with at least one utr | 12486 |
| Number of cds | 22433 |
| Number of exons | 227617 |
| Number of five_prime_utrs | 10044 |
| Number of three_prime_utrs | 5253 |
| Number of exon in cds | 224453 |
| Number of exon in five_prime_utr | 12537 |
| Number of exon in three_prime_utr | 5783 |
| Number of intron in cds | 202020 |
| Number of intron in exon | 205184 |
| Number of intron in five_prime_utr | 2493 |
| Number of intron in three_prime_utr | 530 |
| Number of single exon gene | 91 |
| Number of single exon mrna | 91 |
| mean mrnas per gene | 1 |
| mean cds per mrna | 1 |
| mean exons per mrna | 10.1 |
| mean five_prime_utrs per mrna | 0.4 |
| mean three_prime_utrs per mrna | 0.2 |
| mean exons per cds | 10 |
| mean exons per five_prime_utr | 1.2 |
| mean exons per three_prime_utr | 1.1 |
| mean introns in cds per mrna | 9 |
| mean introns in exons per mrna | 9.1 |
| mean introns in five_prime_utrs per mrna | 0.1 |
| mean introns in three_prime_utrs per mrna | 0 |
| Total gene length | 70550747 |
| Total mrna length | 70550747 |
| Total cds length | 29119128 |
| Total exon length | 30170066 |
| Total five_prime_utr length | 525976 |
| Total three_prime_utr length | 524962 |
| Total intron length per cds | 40117619 |
| Total intron length per exon | 40585865 |
| Total intron length per five_prime_utr | 365056 |
| Total intron length per three_prime_utr | 85815 |
| mean gene length | 3144 |
| mean mrna length | 3144 |
| mean cds length | 1298 |
| mean exon length | 132 |
| mean five_prime_utr length | 52 |
| mean three_prime_utr length | 99 |
| mean cds piece length | 129 |
| mean five_prime_utr piece length | 41 |
| mean three_prime_utr piece length | 90 |
| mean intron in cds length | 198 |
| mean intron in exon length | 197 |
| mean intron in five_prime_utr length | 146 |
| mean intron in three_prime_utr length | 161 |
| Longest genes | 85925 |
| Longest mrnas | 85925 |
| Longest cds | 58605 |
| Longest exons | 36616 |
| Longest five_prime_utrs | 1124 |
| Longest three_prime_utrs | 2255 |
| Longest cds piece | 36616 |
| Longest five_prime_utr piece | 871 |
| Longest three_prime_utr piece | 2230 |
| Longest intron into cds part | 46014 |
| Longest intron into exon part | 46014 |
| Longest intron into five_prime_utr part | 6288 |
| Longest intron into three_prime_utr part | 5958 |
| Shortest genes | 24 |
| Shortest mrnas | 24 |
| Shortest cds | 6 |
| Shortest exons | 2 |
| Shortest five_prime_utrs | 1 |
| Shortest three_prime_utrs | 1 |
| Shortest cds piece | 1 |
| Shortest five_prime_utr piece | 1 |
| Shortest three_prime_utr piece | 1 |
| Shortest intron into cds part | 5 |
| Shortest intron into exon part | 5 |
| Shortest intron into five_prime_utr part | 5 |
| Shortest intron into three_prime_utr part | 5 |
