## Supplementary material for "Phased chromosome-scale genome assembly of an asexual, allopolyploid root-knot nematode reveals complex subgenomic structure: Analysis of the allopolyploid genome of *Meloidogyne javanica*": S4 Fig.docx

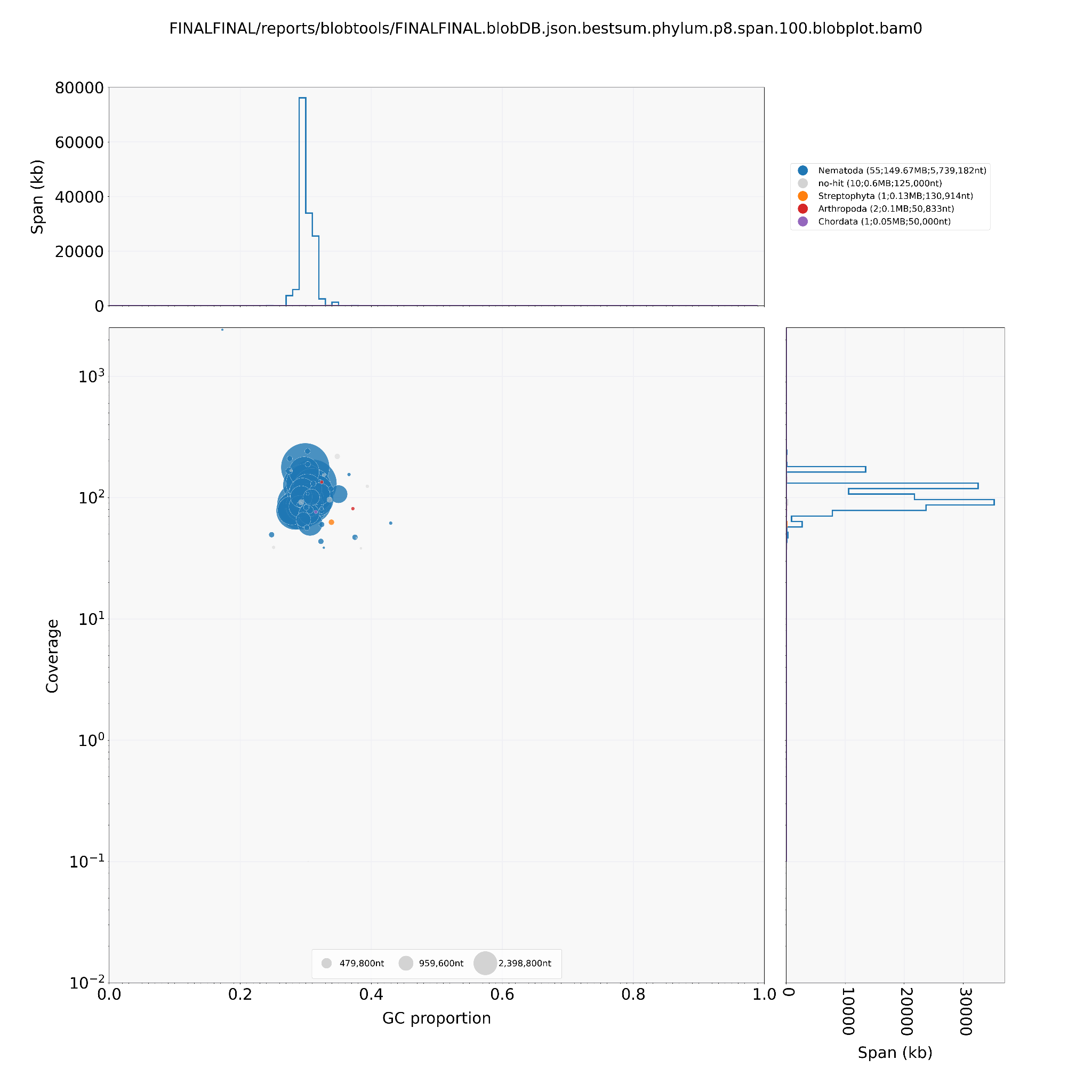

#### Supplementary Figure 4: Blobplot of contamination in our *Meloidogyne javanica* assembly. The x-axis represents the GC proportion, and the y-axis the coverage depth of sequences in the assembly. The size and colour of each blob indicate the relative abundance and taxonomic identity of the contaminating organisms.
