## Supplementary material for "Phased chromosome-scale genome assembly of an asexual, allopolyploid root-knot nematode reveals complex subgenomic structure: Analysis of the allopolyploid genome of *Meloidogyne javanica*": S4 Table.docx

#### Supplementary Table 4: List of pairs that share CDS and amount of shared links.

| **Reference Scaffold** | **Target Scaffold** | **Shared CDS** |
| --- | --- | --- |
| 2 | 13 | 46 |
| 3 | 11 | 284 |
| 4 | 14 | 307 |
| 5 | 12 | 182 |
| 6 | 19 | 216 |
| 6 | 1 | 206 |
| 7 | 10 | 206 |
| 8 | 16 | 273 |
| 8 | 33 | 162 |
| 8 | 29 | 58 |
| 8 | 28 | 49 |
| 9 | 5 | 102 |
| 13 | 15 | 351 |
| 17 | 25 | 112 |
| 18 | 26 | 63 |
| 20 | 23 | 214 |
| 20 | 32 | 120 |
| 24 | 30 | 20 |
| 25 | 27 | 64 |
