## Supplementary material for "Phased chromosome-scale genome assembly of an asexual, allopolyploid root-knot nematode reveals complex subgenomic structure: Analysis of the allopolyploid genome of *Meloidogyne javanica*": S5 Fig.docx

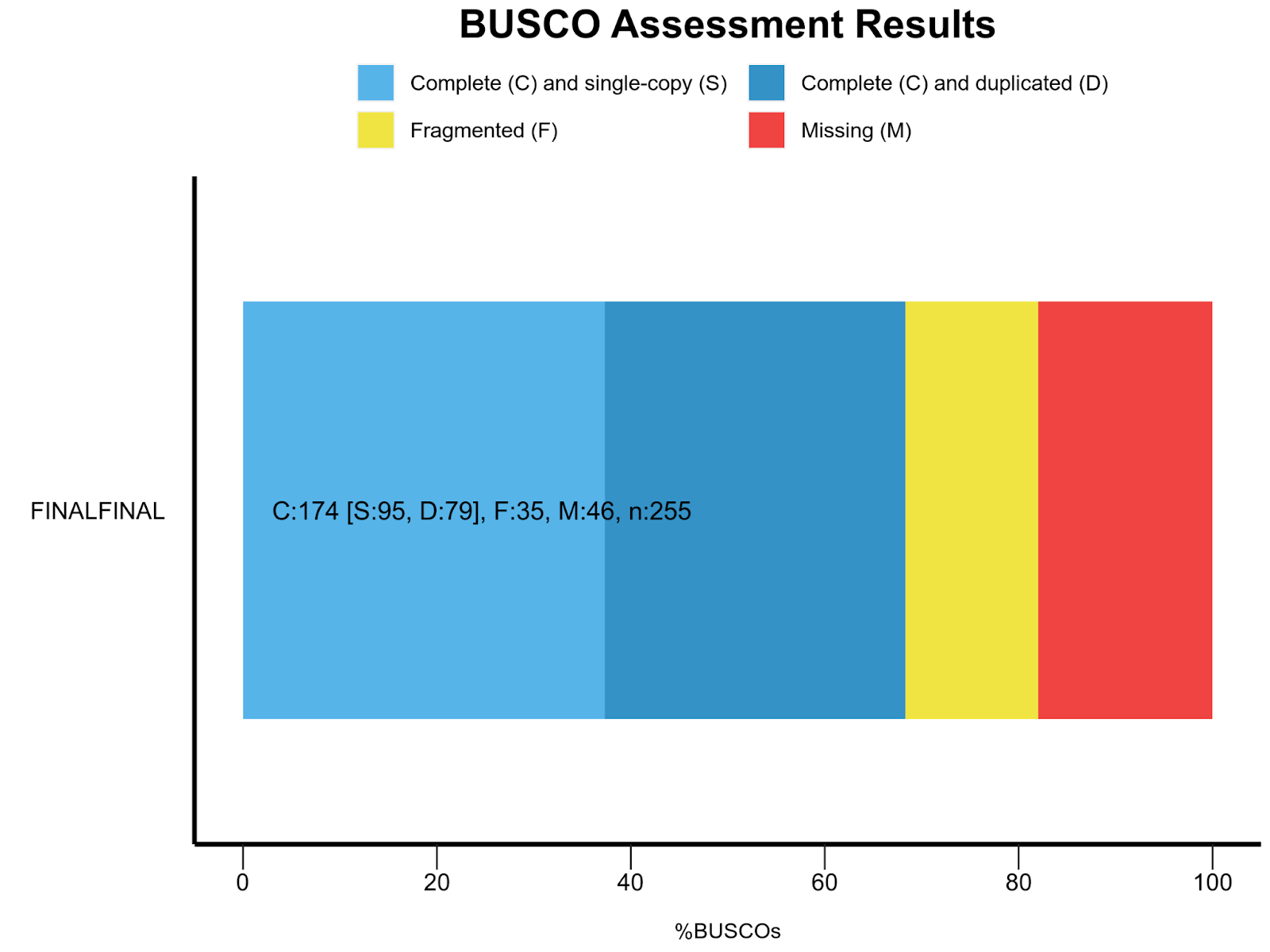

#### Supplementary Figure 5: BUSCO v5 results. Barplot showing the proportion of BUSCO genes identified in the assembly. Analysis performed using the *eukaryota_odb10* database. Generated by *BUSCO.*
