## Supplementary material for "Phased chromosome-scale genome assembly of an asexual, allopolyploid root-knot nematode reveals complex subgenomic structure: Analysis of the allopolyploid genome of *Meloidogyne javanica*": S5 Table.docx

#### Supplementary Table 5: Validation of homoeologous pairings.

| **Scaffold number** | **Orthology** | **MASH** | **BUSCO** | **Consensus homoeologous counterpart** |
| --- | --- | --- | --- | --- |
| 1 | 6 | 6 | 6 | 6 |
| 2 | 13 | - | 52 | 13 |
| 3 | 11 | 11 | 7 | 11 |
| 4 | 14 | 14 | 6 | 14 |
| 5 | 12 | 12 | 9 | 12 |
| 6 | 21 | 21 | - | 21 |
| 7 | 10 | 10 | - | 10 |
| 8 | 16 | 16 | - | 16 |
| 9 | 5 | - | 7 | 5 |
| 10 | 7 | 7 | 3 | 7 |
| 11 | 3 | 3 | 5 | 3 |
| 12 | 5 | 5 | 15 | 5 |
| 13 | 15 | 15 | 45 | 15 |
| 14 | 4 | 4 | - | 4 |
| 15 | 13 | 13 | 8 | 13 |
| 16 | 8 | 8 | 25 | 8 |
| 17 | 25 | 25 | 26 | 25 |
| 18 | 26 | - | 21 | 26 |
| 19 | 6 | 6 | 30 | 6 |
| 20 | 23 | 23 | 6 | 23 |
| 21 | 6 | 6 | - | 6 |
| 22 | 10 | 10 | - | 10 |
| 23 | 20 | 20 | - | 20 |
| 24 | 30 | - | - | 30 |
| 25 | 17 | 27 | - | 17 |
| 26 | 18 | - | 49 | 18 |
| 27 | 25 | - | - | 25 |
| 28 | 8 | - | 8 | 8 |
| 29 | 8 | - | - | 8 |
| 30 | 24 | - | - | 24 |
| 31 | 6 | - | - | 6 |
| 32 | 20 | - | - | 20 |
| 33 | 8 | - | - | 8 |
