## Supplementary material for "Phased chromosome-scale genome assembly of an asexual, allopolyploid root-knot nematode reveals complex subgenomic structure: Analysis of the allopolyploid genome of *Meloidogyne javanica*": S6 Fig.docx

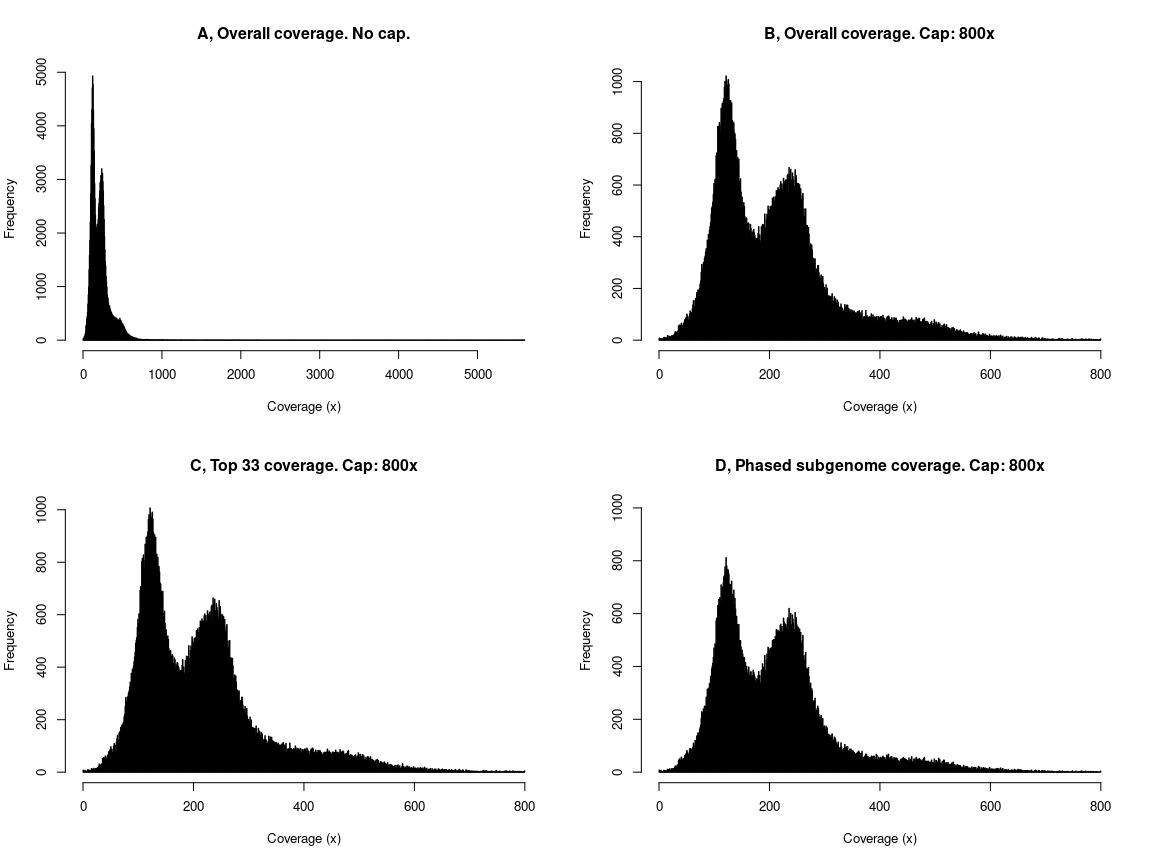

#### Supplementary Figure 6: Assembly-wide distributions of coverage depth. A, Distribution of coverage depth at all bases in the assembly. B, Distribution of coverage depth of all bases in the assembly, capped at 800x. C, Distribution of coverage of all bases in the longest 33 scaffolds, capped at 800x. D, Distribution of coverage of all bases in scaffolds phased to a subgenome, capped at 800x.
