## Supplementary material for "Phased chromosome-scale genome assembly of an asexual, allopolyploid root-knot nematode reveals complex subgenomic structure: Analysis of the allopolyploid genome of *Meloidogyne javanica*": S6 Table.docx

#### Supplementary Table 6: Summary of longest 33 scaffolds of diploid assembly of *M. javanica*.

| **Scaffold Number** | **Length (bp)** | **Mean coverage (x)** | **Phase assignment (A, B, or unphased)** | **Comments** |
| --- | --- | --- | --- | --- |
| 1 | 9595054 | 245 | B | Some collapse. |
| 2 | 9577269 | 337 | U | Extensive collapse. |
| 3 | 8254935 | 181 | A |  |
| 4 | 7520906 | 251 | A | Some collapse. |
| 5 | 7228418 | 184 | A |  |
| 6 | 7199321 | 177 | A |  |
| 7 | 7044505 | 169 | A |  |
| 8 | 6281301 | 186 | A |  |
| 9 | 6143287 | 260 | B | Some collapse. |
| 10 | 5957753 | 164 | B |  |
| 11 | 5739182 | 247 | B |  |
| 12 | 5730042 | 205 | B |  |
| 13 | 5673700 | 219 | B |  |
| 14 | 4766193 | 189 | B |  |
| 15 | 4415488 | 196 | A |  |
| 16 | 4345108 | 204 | B | Some collapse. |
| 17 | 3893252 | 201 | B |  |
| 18 | 3682237 | 314 | B | Some collapse. |
| 19 | 3649086 | 168 | U | Some collapse. |
| 20 | 3413746 | 207 | A |  |
| 21 | 3407168 | 264 | B |  |
| 22 | 3386470 | 162 | A | Low coverage, possibly single copy. Phased manually. |
| 23 | 2744999 | 231 | B | Phased manually. |
| 24 | 2512086 | 131 | U | Low coverage, possibly single copy. |
| 25 | 2178917 | 178 | A |  |
| 26 | 2011659 | 171 | A |  |
| 27 | 2002407 | 183 | B | Phased manually. |
| 28 | 1997981 | 214 | B | Phased manually. |
| 29 | 1951323 | 210 | B |  |
| 30 | 1735511 | 151 | U |  |
| 31 | 1316575 | 219 | U |  |
| 32 | 1091457 | 211 | B |  |
| 33 | 898067 | 125 | B | Low coverage, possibly single copy. Phased manually. |

^1^ Here collapse refers to regions of a scaffold with more than 2 copies predicted based on read depth.
