## Supplementary material for "Phased chromosome-scale genome assembly of an asexual, allopolyploid root-knot nematode reveals complex subgenomic structure: Analysis of the allopolyploid genome of *Meloidogyne javanica*": S7 Fig.docx

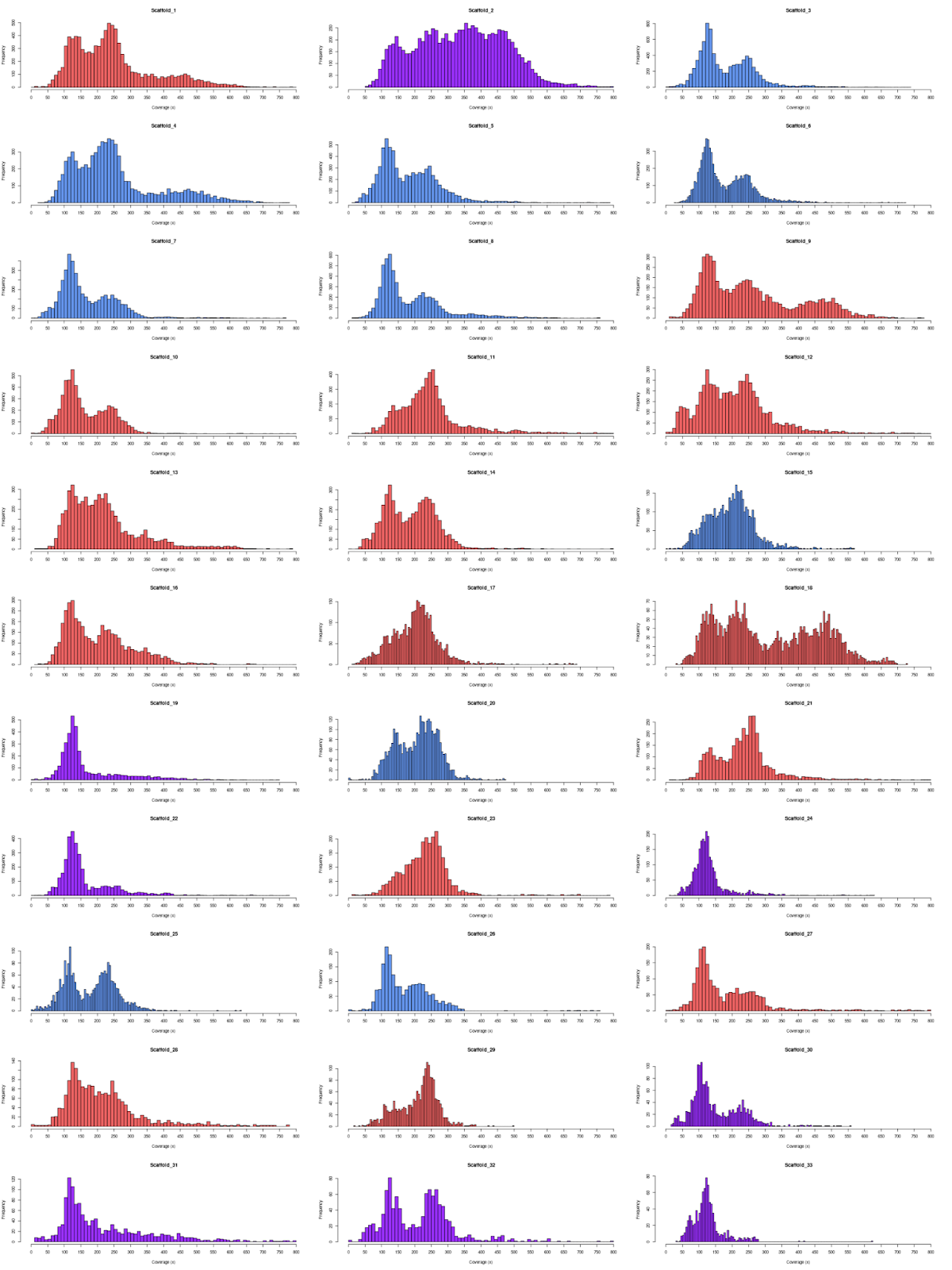

#### Supplementary Figure 7: Scaffold-level distributions of coverage depth. Subgenome A in blue, subgenome B in red, and unphased scaffolds in purple. X-axis shows coverage depth, y-axis shows frequency.
