## Supplementary material for "Phased chromosome-scale genome assembly of an asexual, allopolyploid root-knot nematode reveals complex subgenomic structure: Analysis of the allopolyploid genome of *Meloidogyne javanica*": S8 Fig.docx

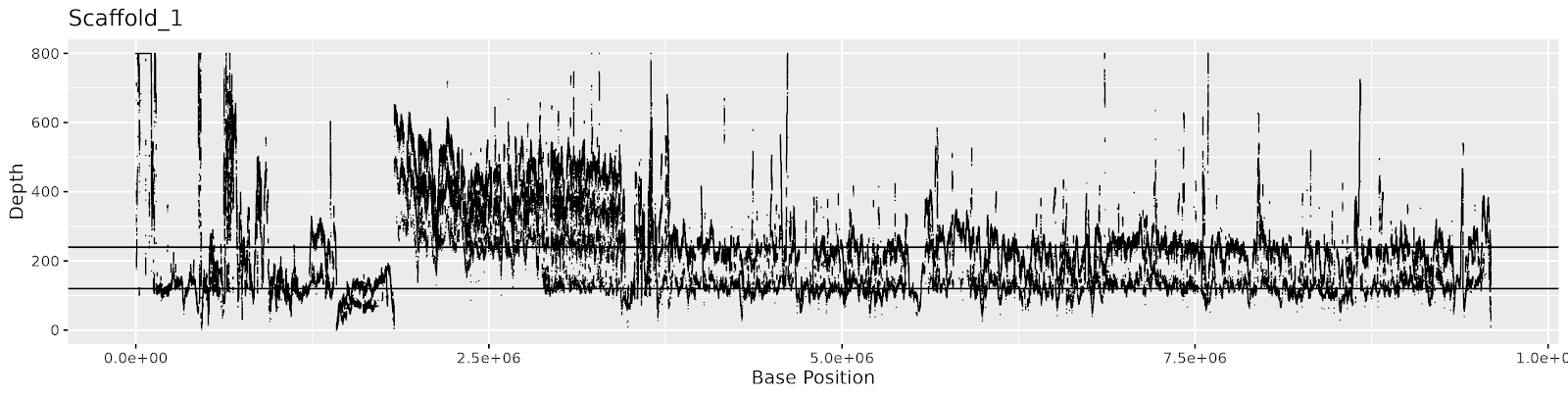

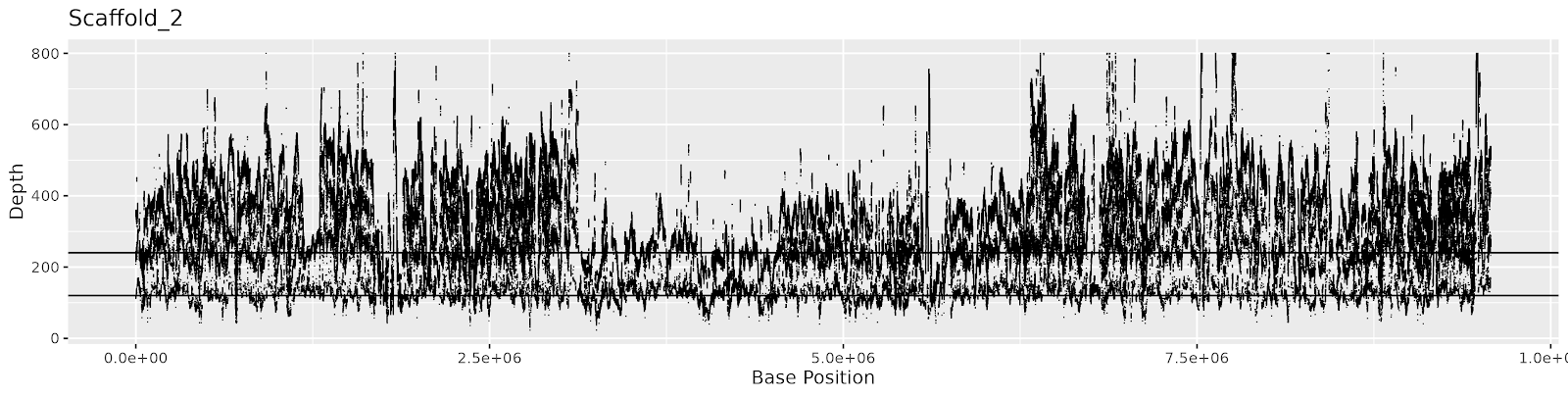

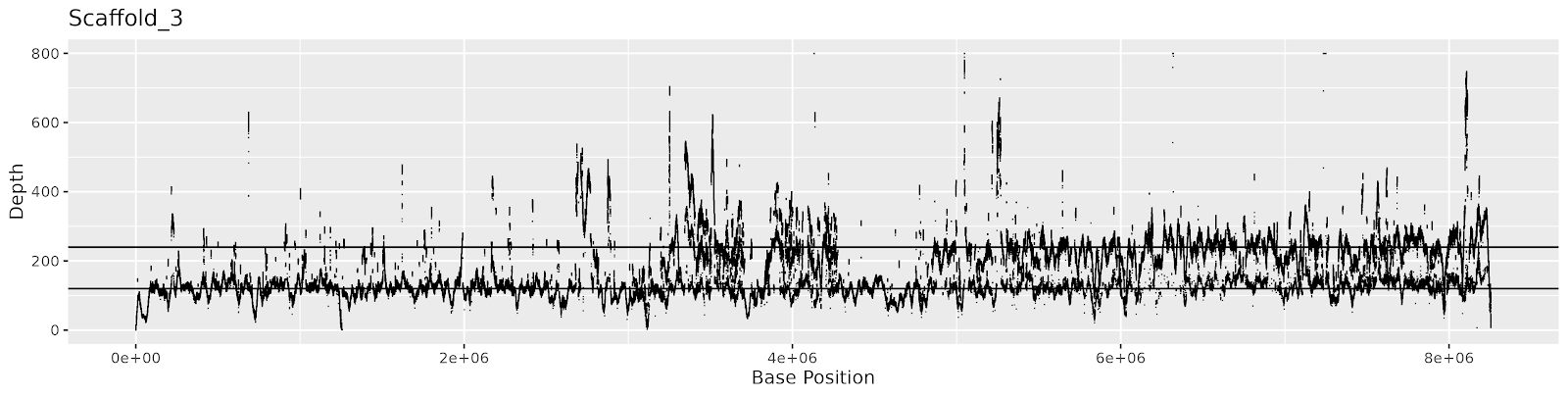

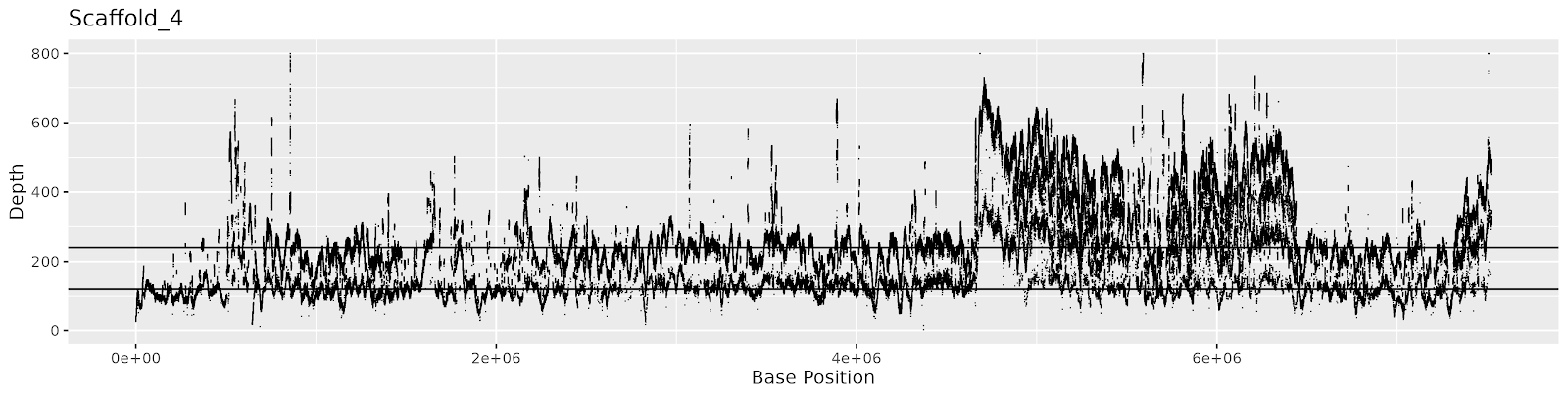

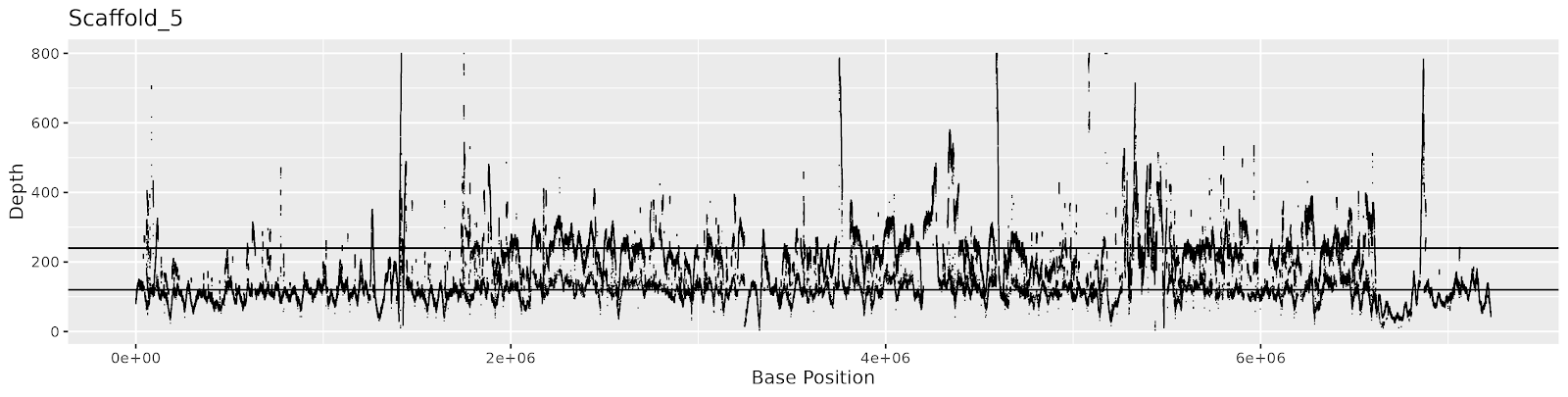

#### Supplementary Figure 8a: Coverage depth of individual bases across scaffolds 1-5. Horizontal lines mark 120x and 240x coverage depth, corresponding to peaks seen in Supplementary Figure 7.

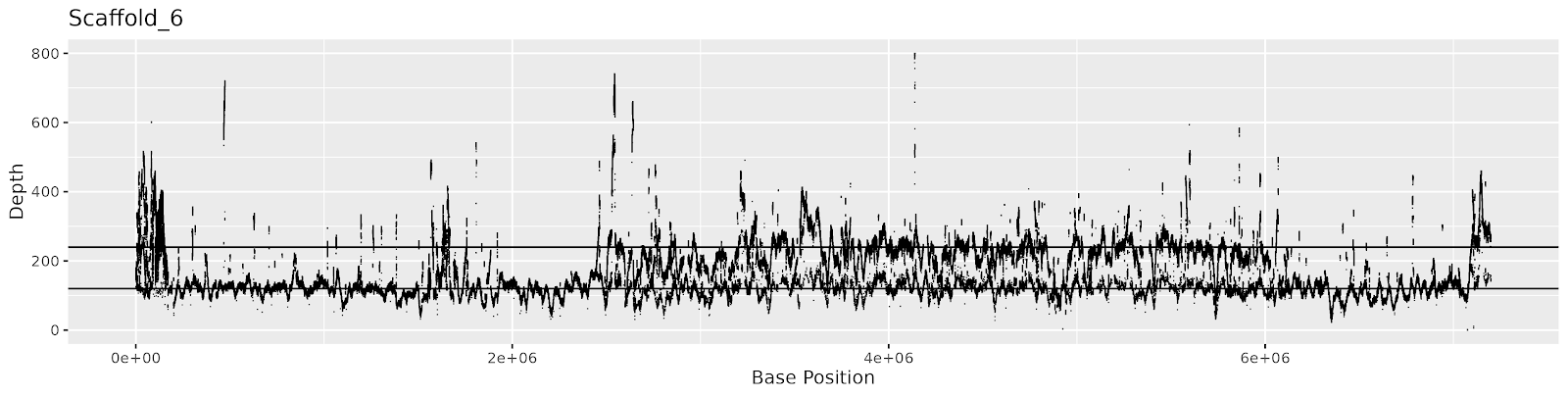

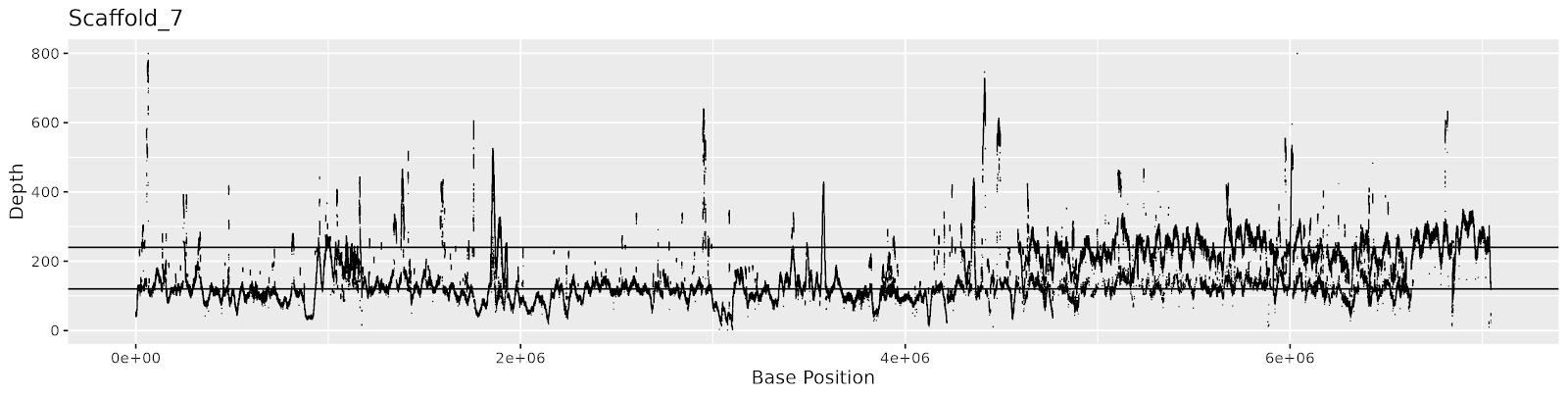

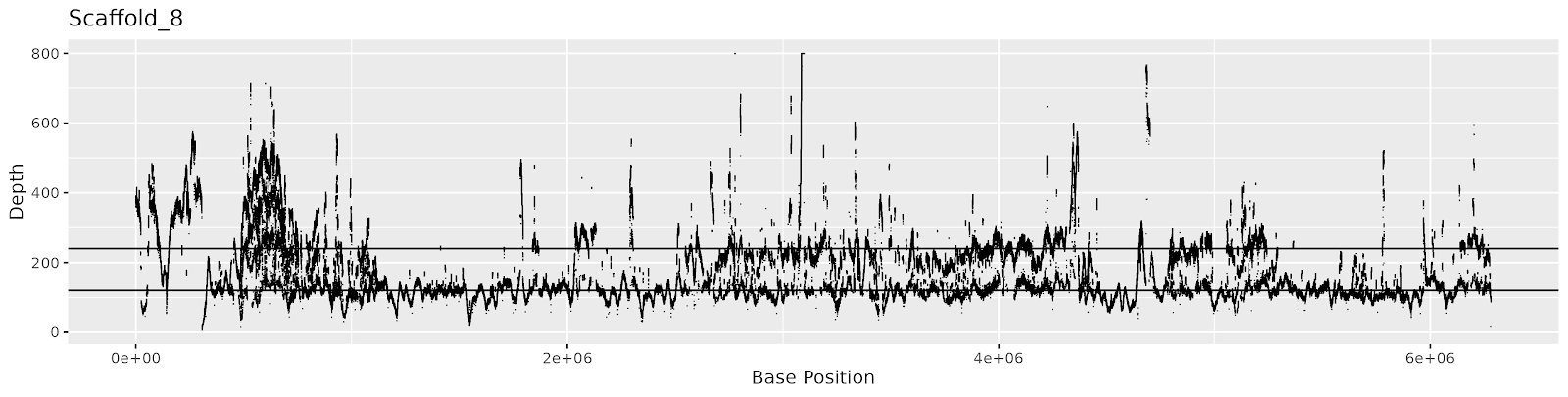

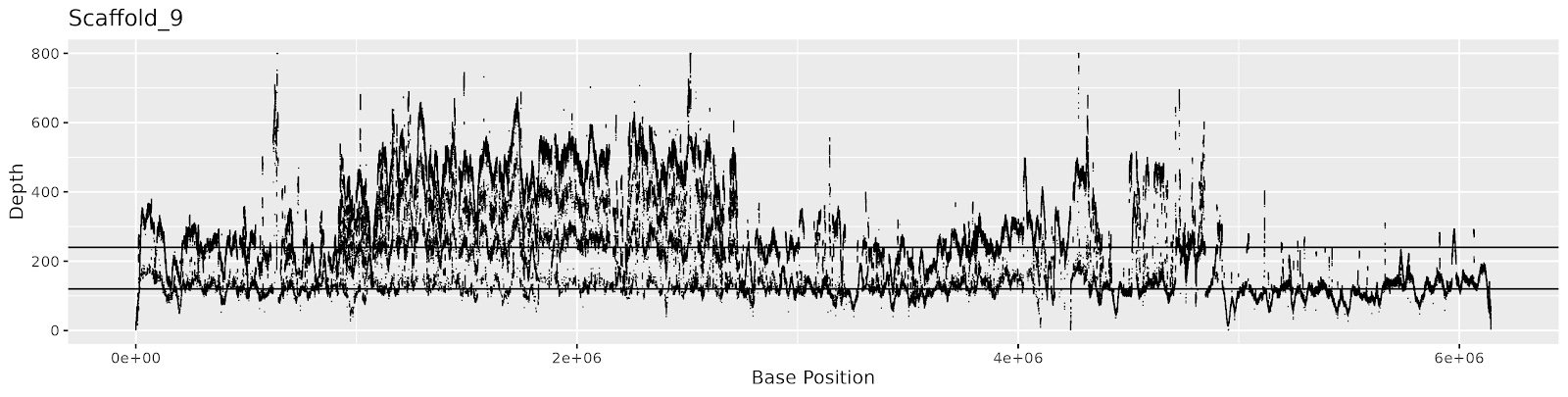

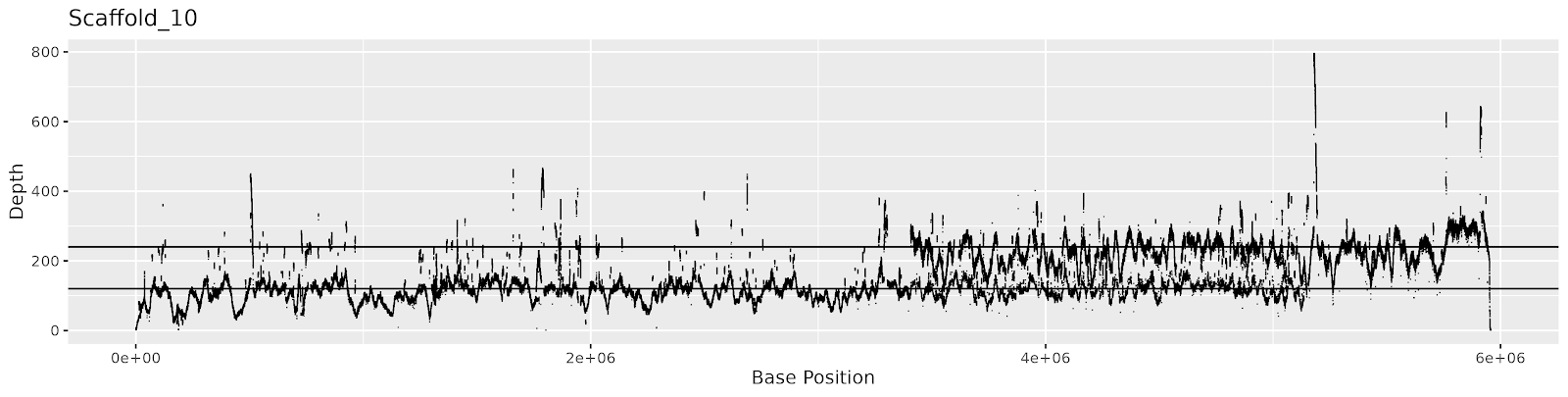

#### Supplementary Figure 8b: Coverage depth of individual bases across scaffolds 6-10. Horizontal lines mark 120x and 240x coverage depth, corresponding to peaks seen in Supplementary Figure 7.

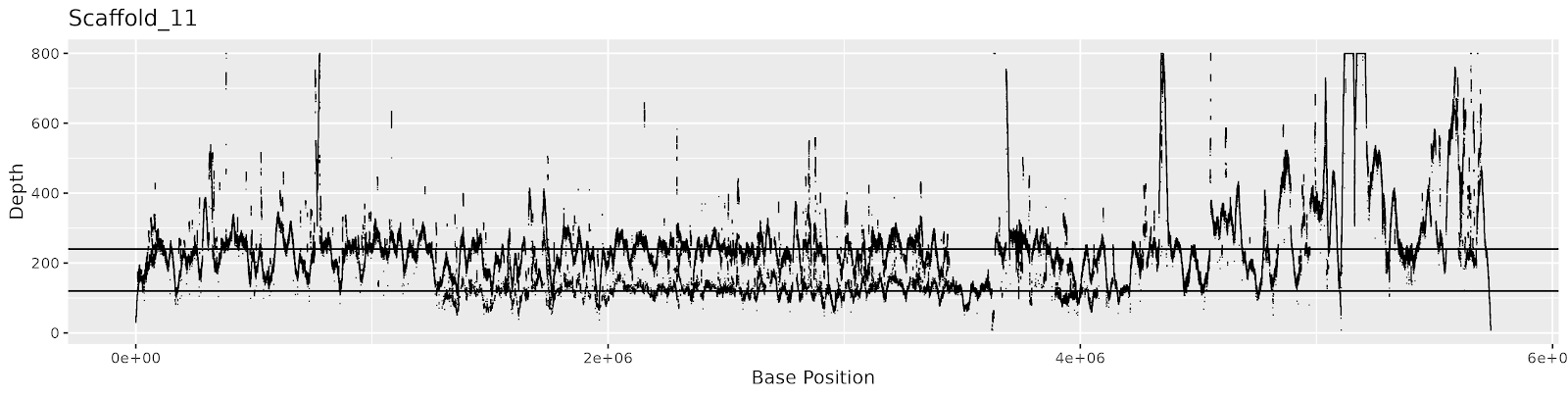

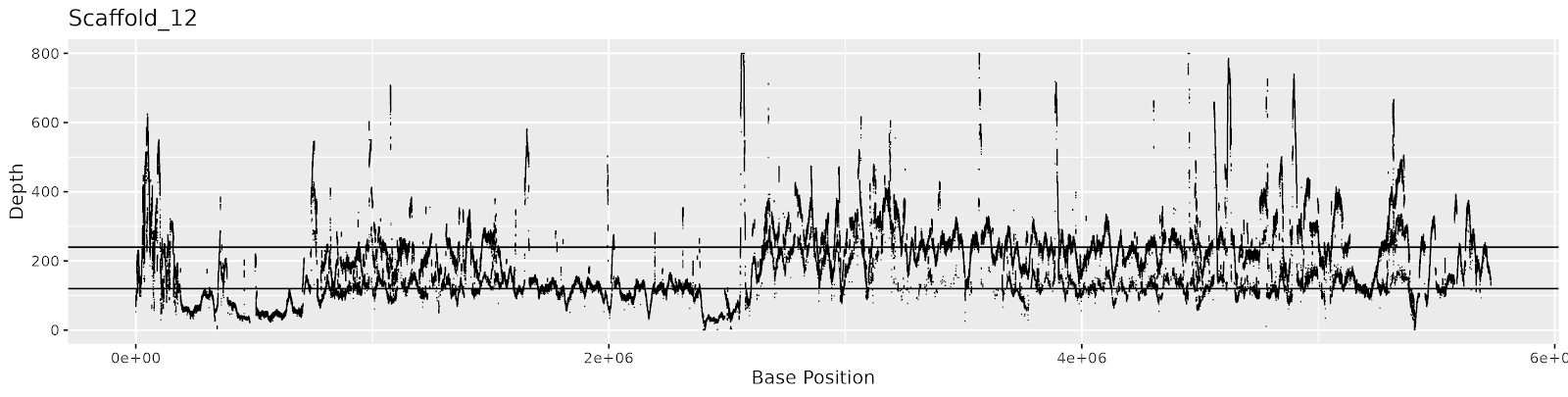

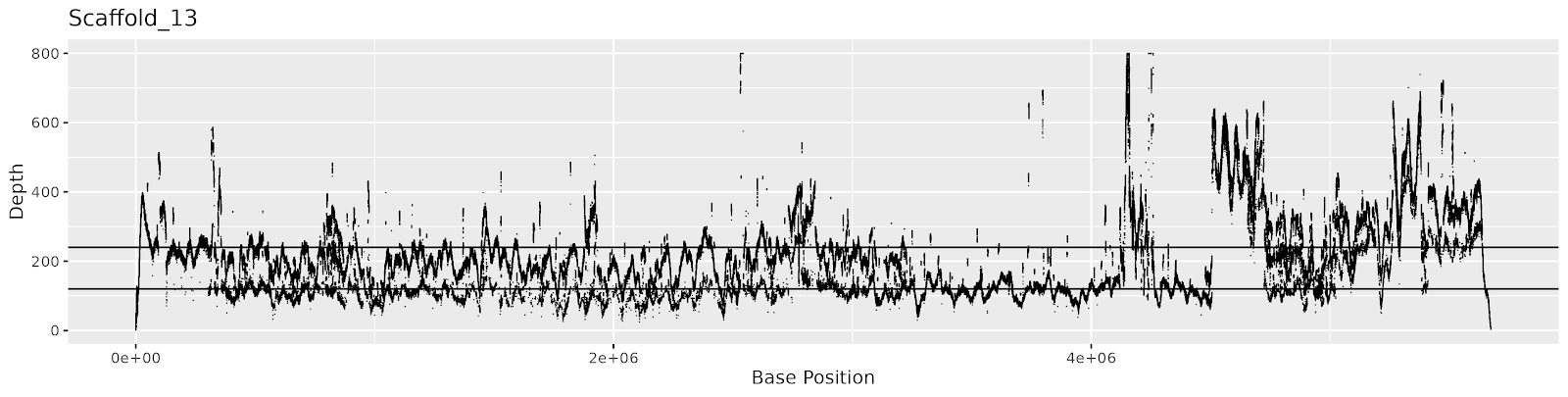

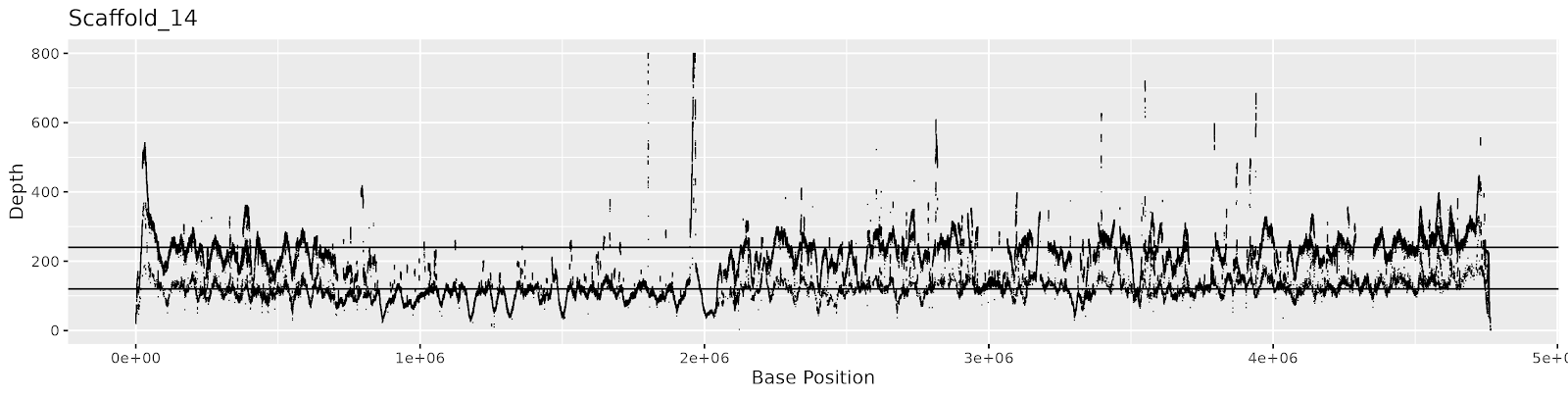

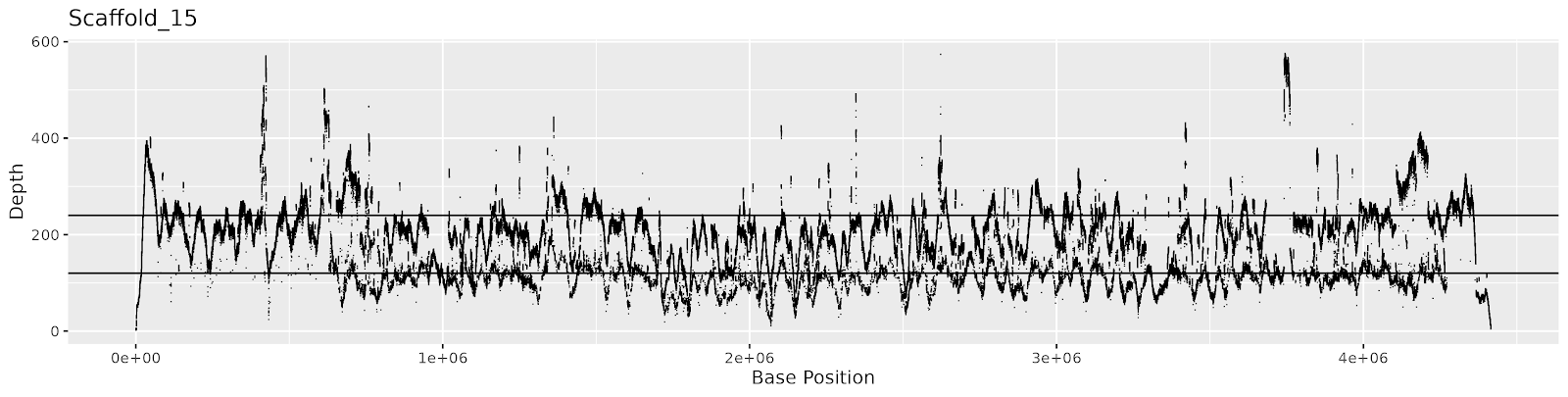

#### Supplementary Figure 8c: Coverage depth of individual bases across scaffolds 11-15. Horizontal lines mark 120x and 240x coverage depth, corresponding to peaks seen in Supplementary Figure 7.

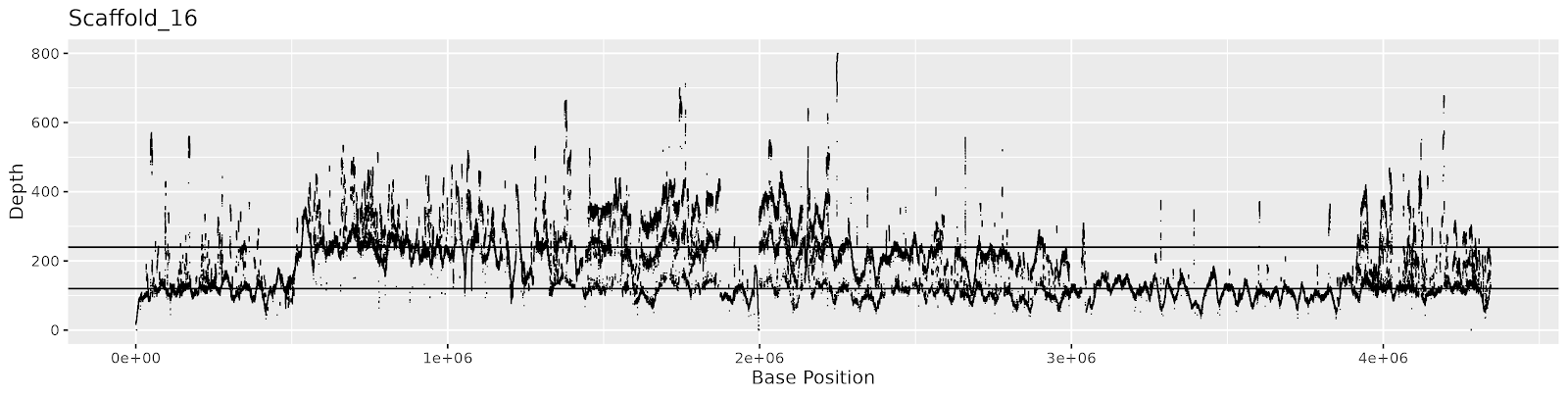

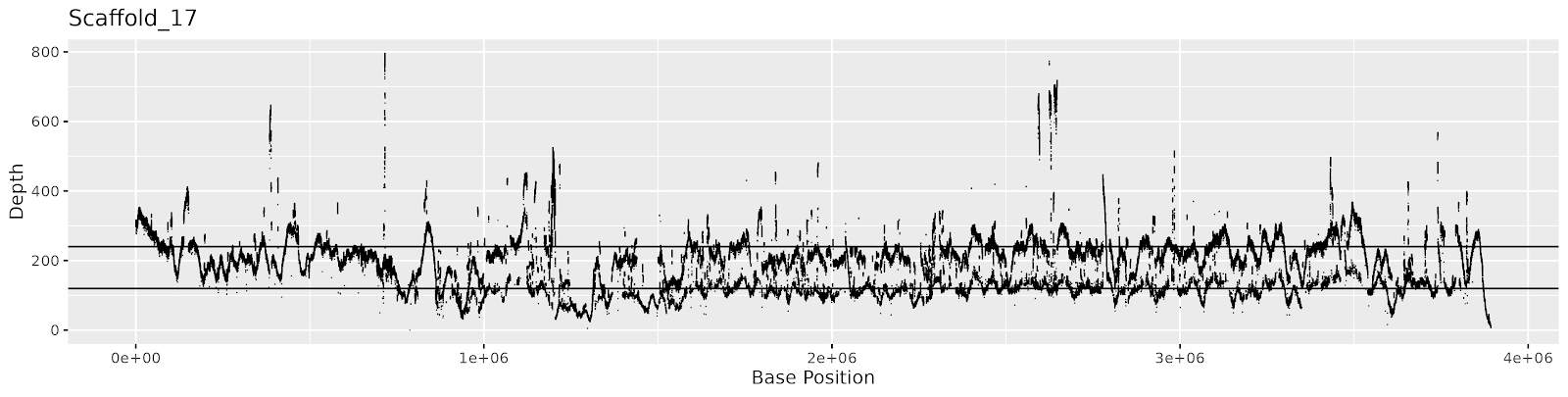

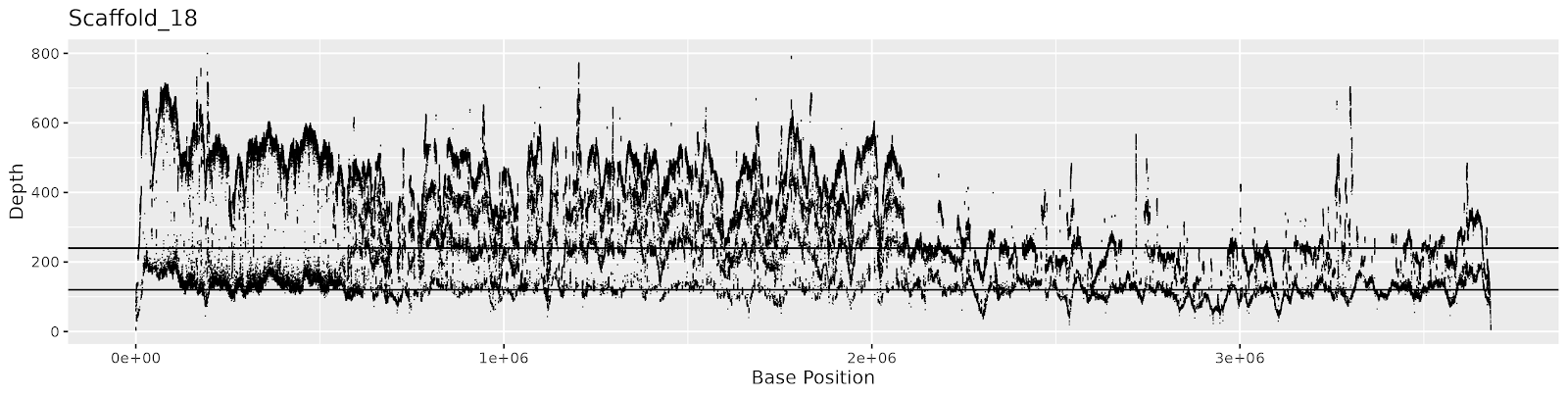

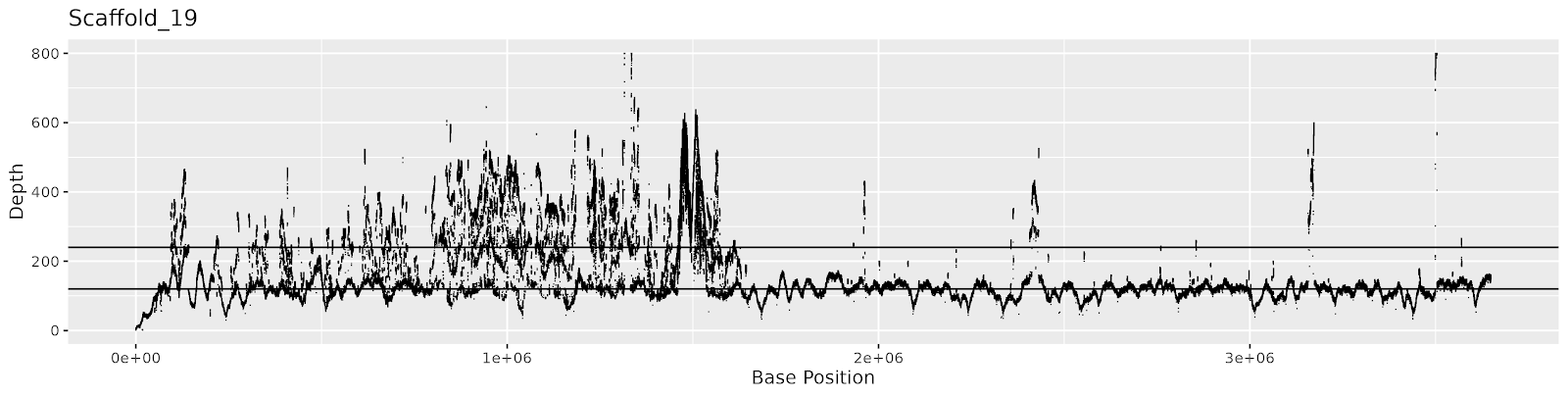

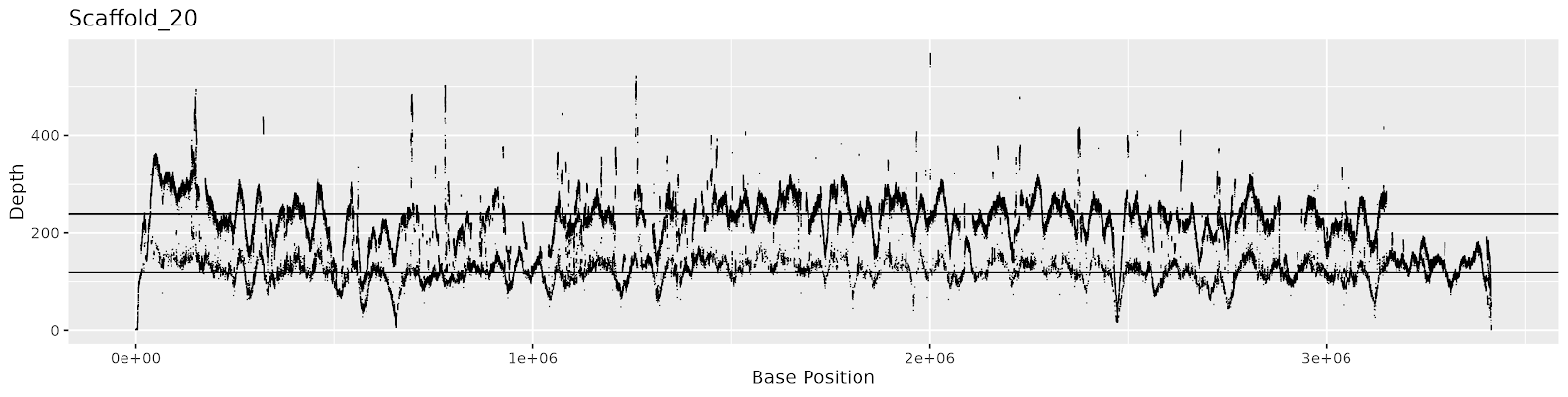

#### Supplementary Figure 8d: Coverage depth of individual bases across scaffolds 16-20. Horizontal lines mark 120x and 240x coverage depth, corresponding to peaks seen in Supplementary Figure 7.

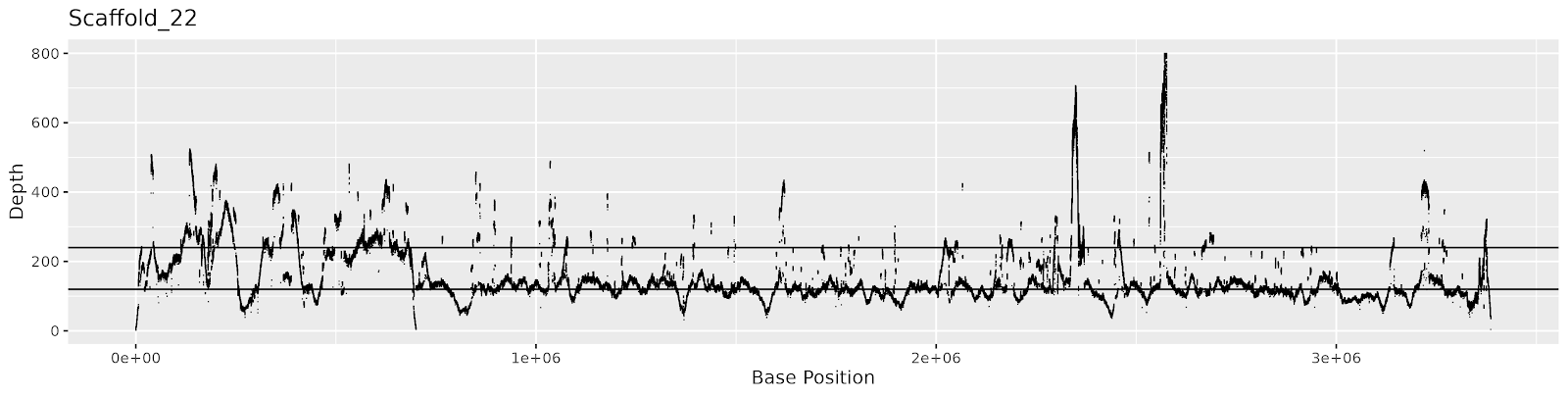

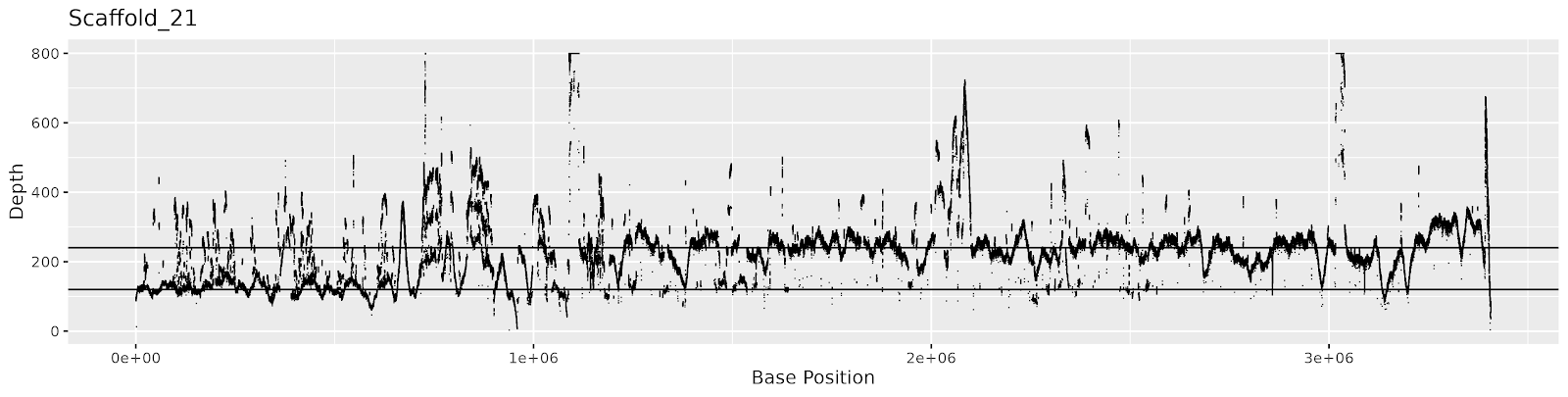

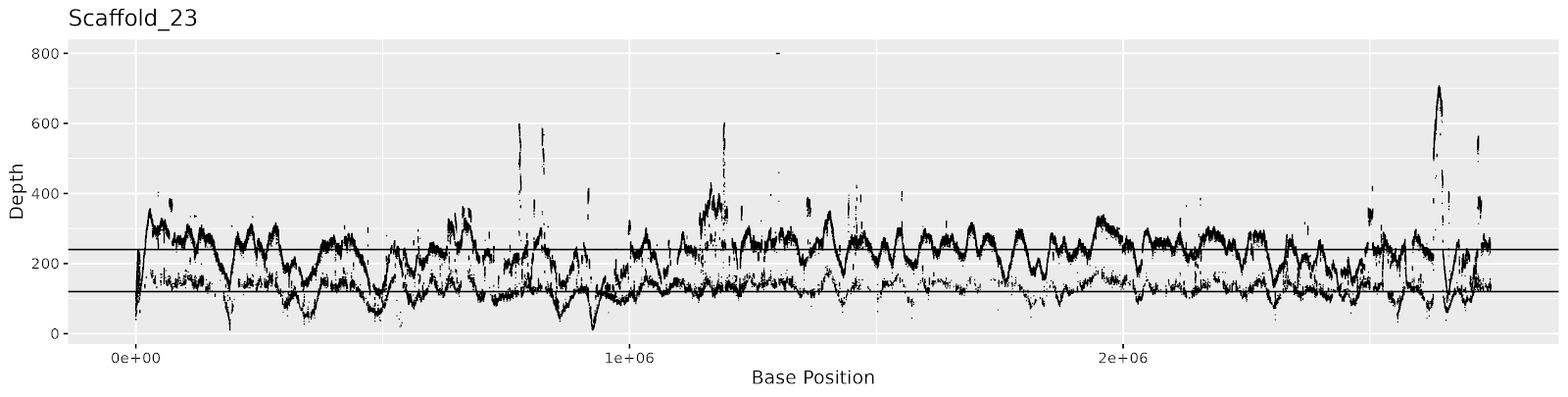

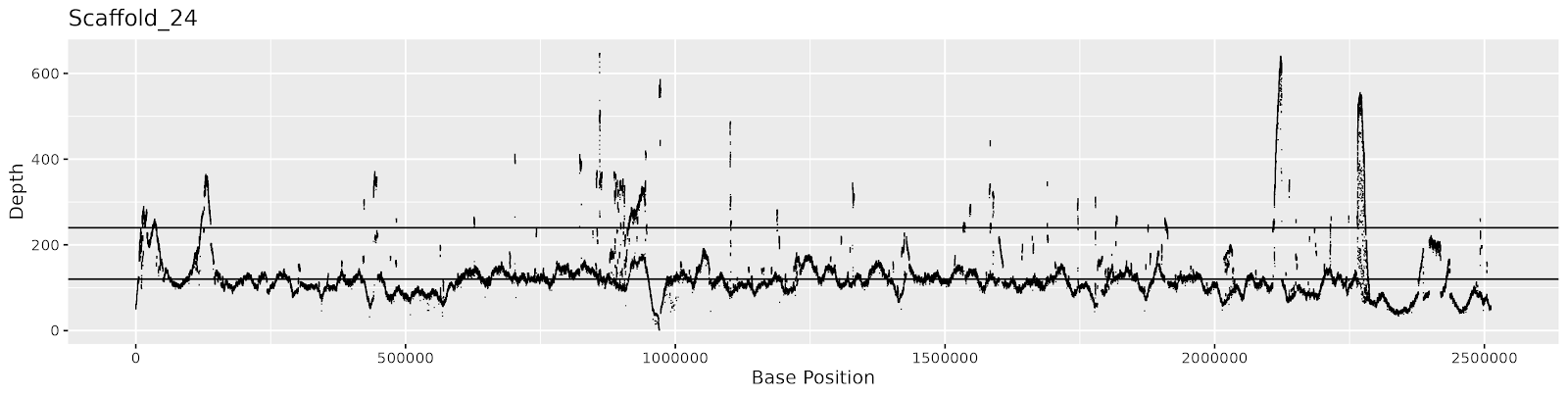

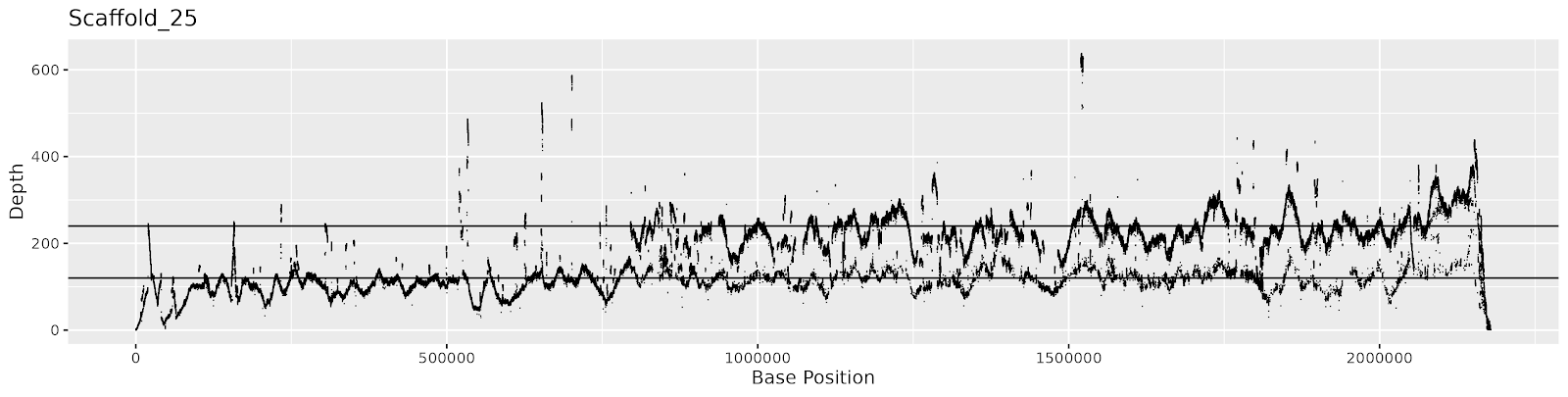

#### Supplementary Figure 8d: Coverage depth of individual bases across scaffolds 21-25. Horizontal lines mark 120x and 240x coverage depth, corresponding to peaks seen in Supplementary Figure 7.

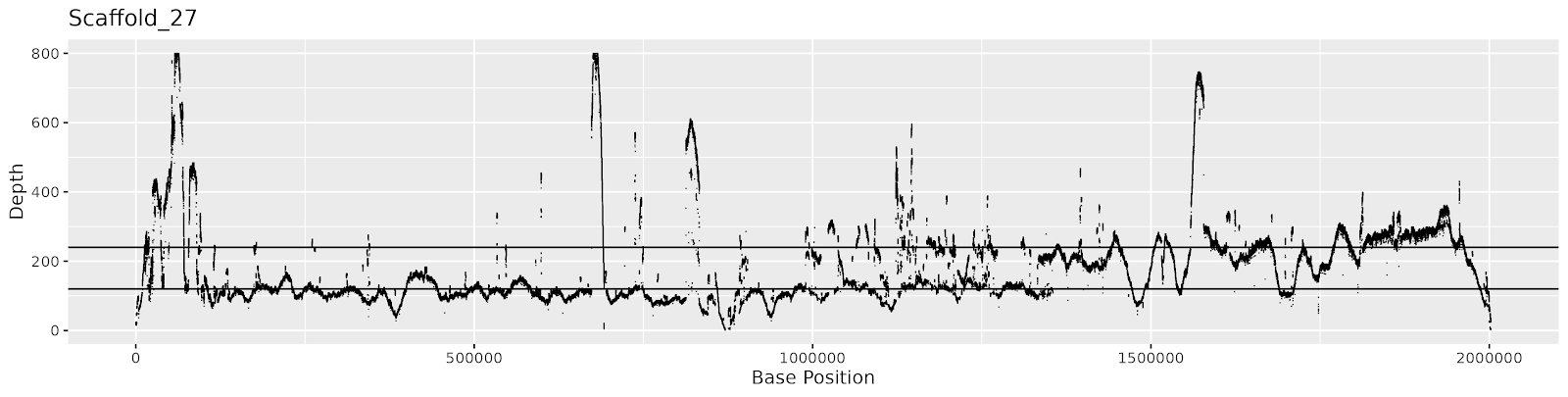

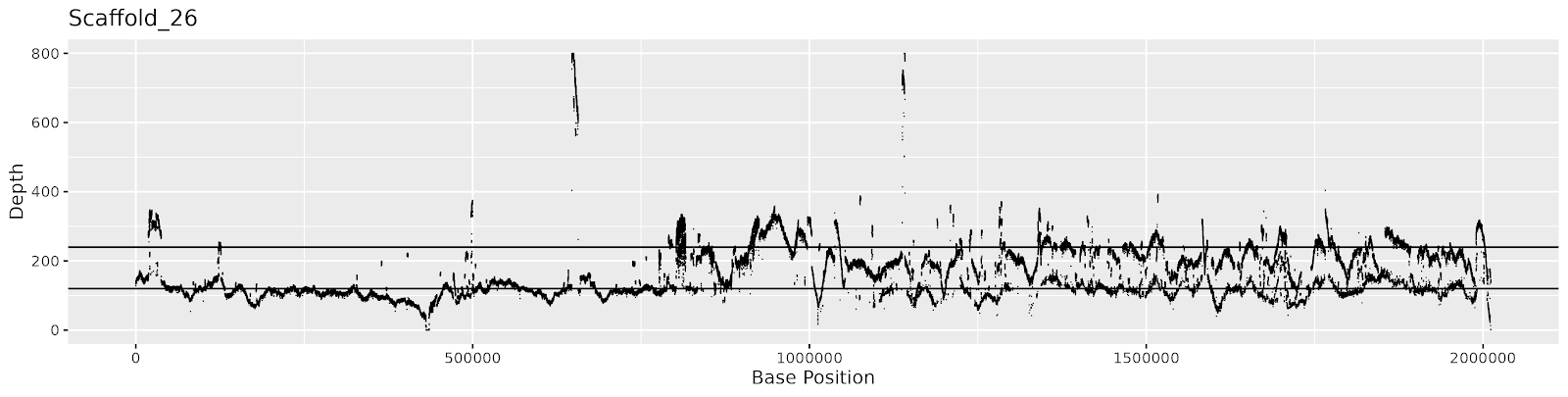

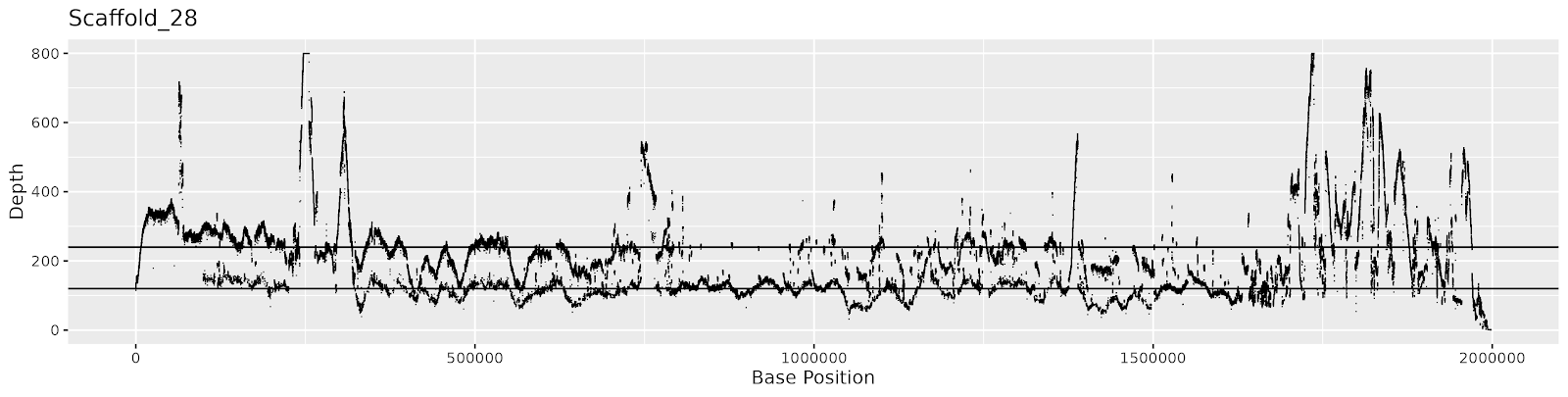

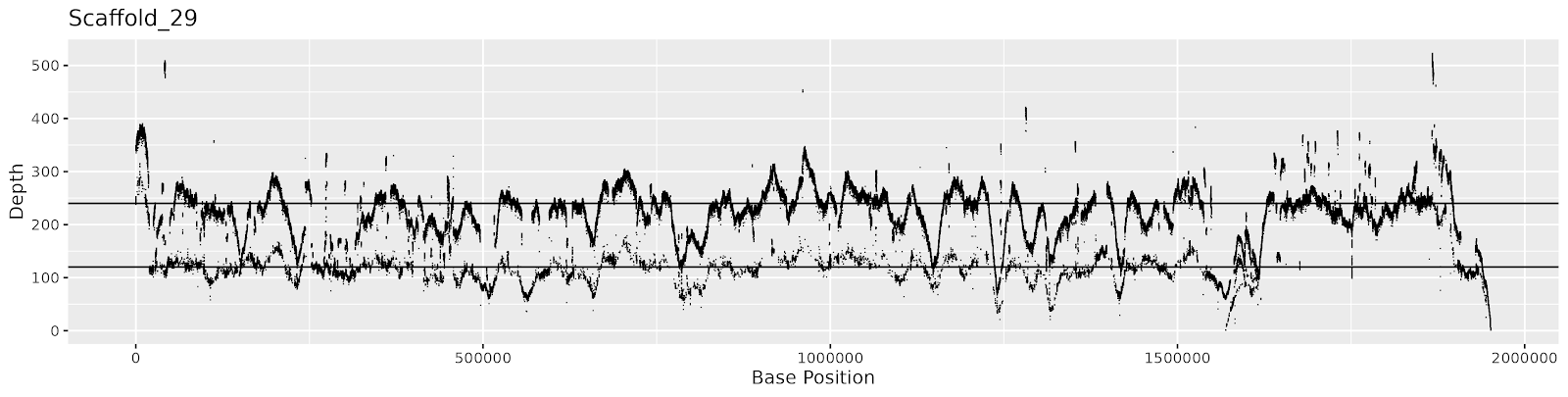

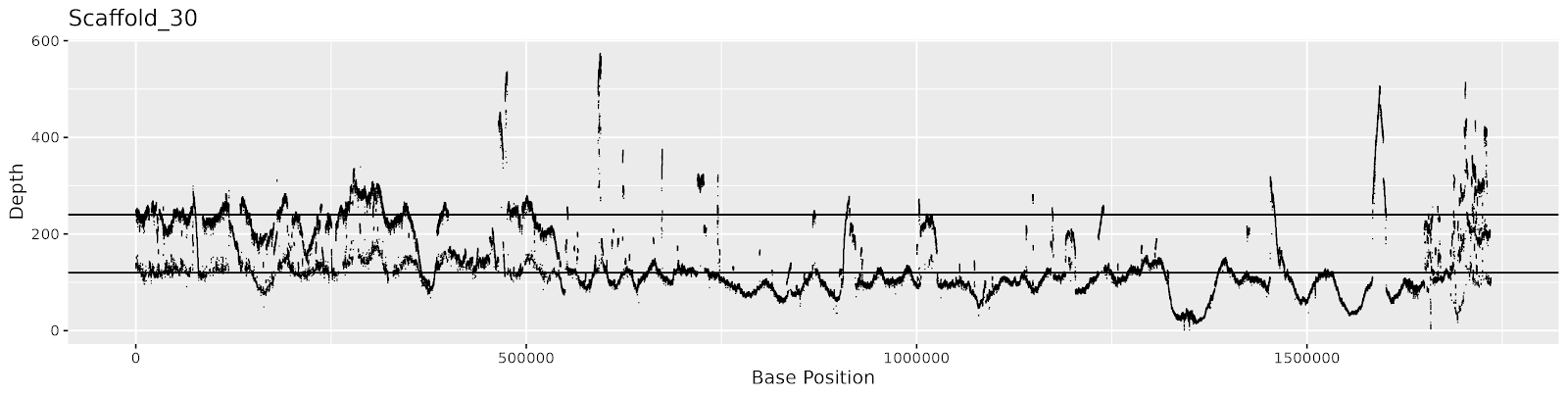

#### Supplementary Figure 8e: Coverage depth of individual bases across scaffolds 26-30. Horizontal lines mark 120x and 240x coverage depth, corresponding to peaks seen in Supplementary Figure 7.

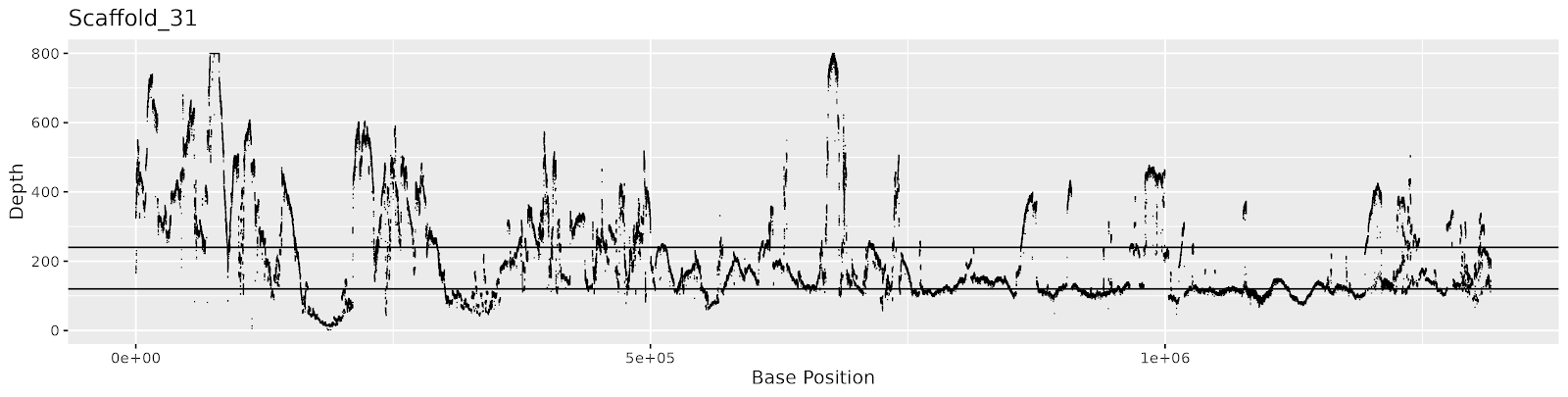

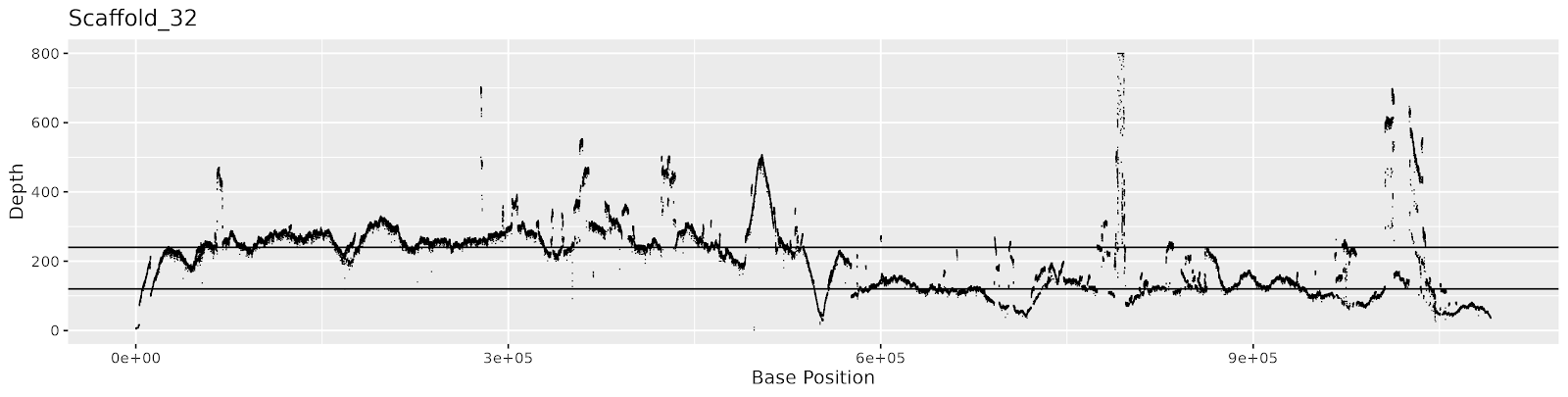

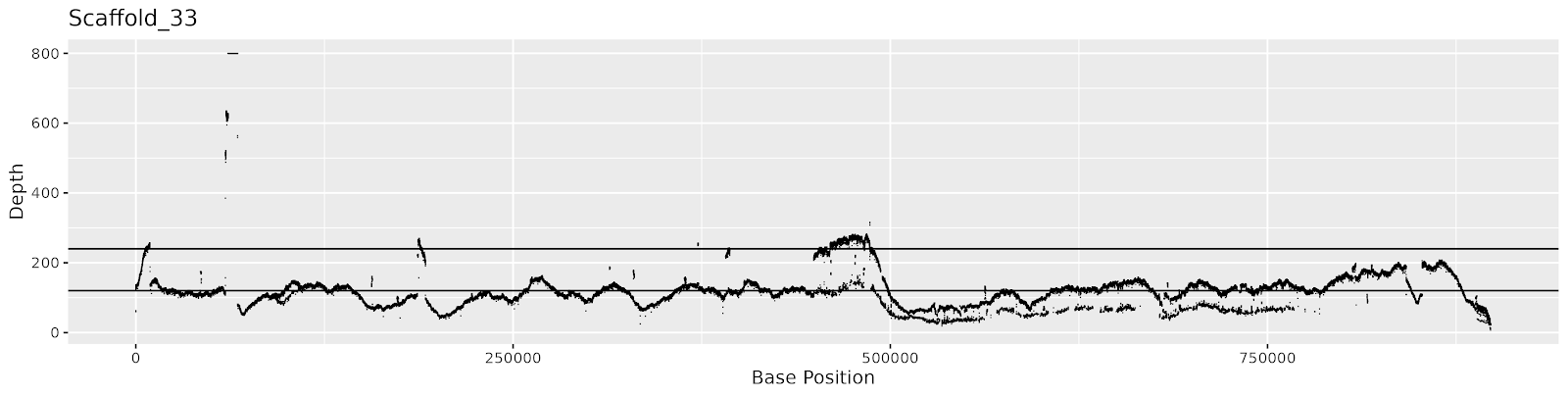

#### Supplementary Figure 8f: Coverage depth of individual bases across scaffolds 31-33. Horizontal lines mark 120x and 240x coverage depth, corresponding to peaks seen in Supplementary Figure 7.
