## Supplementary figures and images for "Phased chromosome-scale genome assembly of an asexual, allopolyploid root-knot nematode reveals complex subgenomic structure: Analysis of the allopolyploid genome of *Meloidogyne javanica*"

### S9 Fig.docx

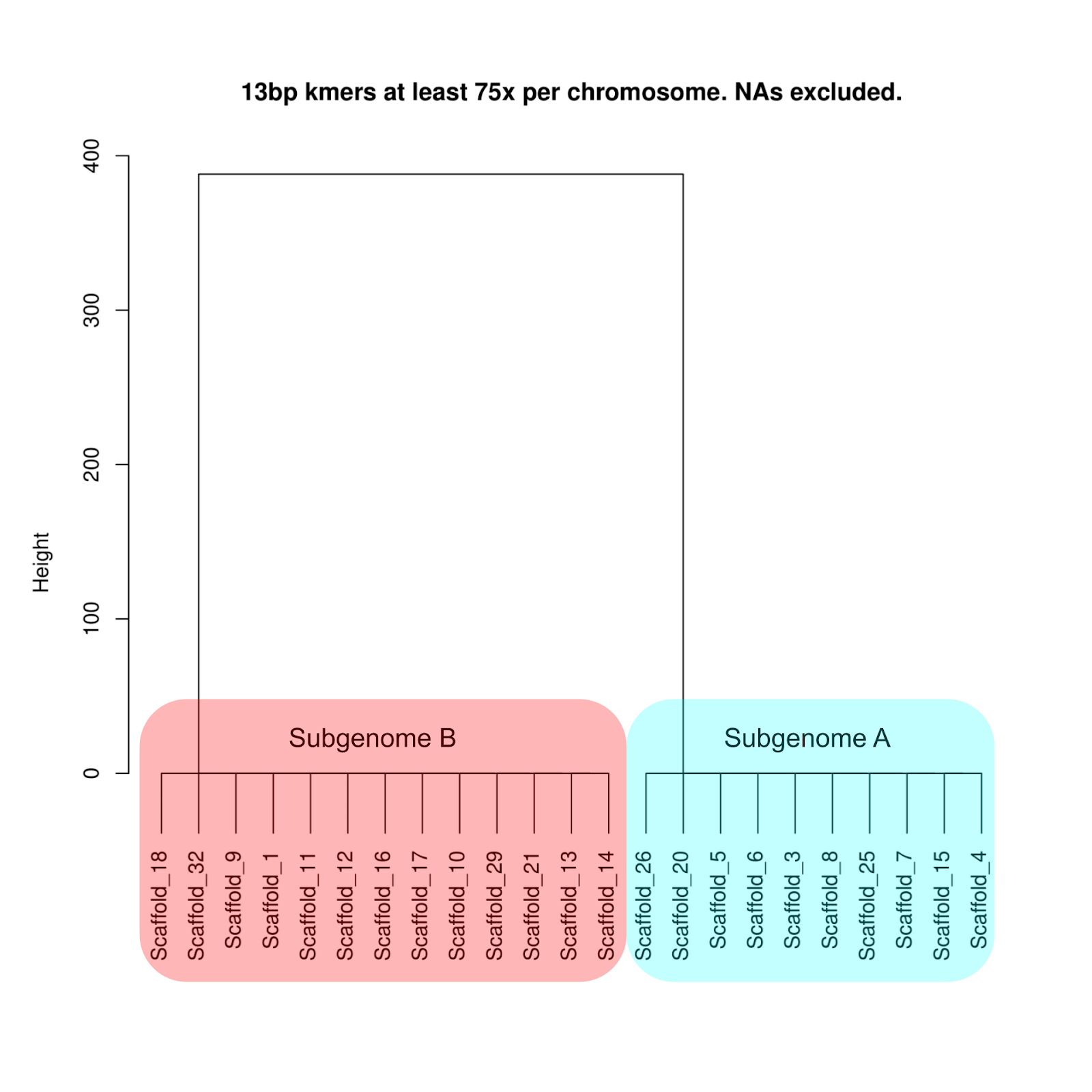

#### Supplementary Figure 9: Subgenome-phased dendrogram of scaffolds.
